# Nicotinic cholinergic receptors in VTA glutamate neurons modulate excitatory transmission

**DOI:** 10.1101/271205

**Authors:** Yijin Yan, Can Peng, Matthew C. Arvin, Xiao-Tao Jin, Veronica J. Kim, Matthew D. Ramsey, Yong Wang, Sambashiva Banala, David L. Wokosin, J. Michael McIntosh, Luke D. Lavis, Luke D. Lavis, Ryan M. Drenan

## Abstract

Ventral tegmental area (VTA) glutamate neurons are important components of brain reward circuitry, but whether they are subject to cholinergic modulation is unknown. To study this, we used an array of molecular, physiological, and photostimulation techniques to examine nicotinic acetylcholine receptors (nAChRs) in VTA glutamate neurons. VTA neurons positive for the vesicular glutamate transporter 2 (VGLUT2+) are responsive to acetylcholine (ACh) released from mesopontine cholinergic axons. VTA VGLUT2+ neurons express mRNA and protein subunits known to comprise typical heteromeric nAChRs. Electrophysiology, coupled with 2-photon microscopy and laser flash photolysis of a photoactivatable nicotine probe, was used to demonstrate nAChR functional activity in the somatodendritic subcellular compartment of VTA VGLUT2+ neurons. Finally, optogenetic isolation of intrinsic VTA glutamatergic microcircuits demonstrated that nicotine potently modulates excitatory transmission within the VTA. These results indicate that VTA glutamate neurons are modulated by cholinergic mechanisms and participate in the cascade of physiological responses to nicotine exposure.

## Introduction

Nicotine, the primary psychoactive agent in tobacco products, activates and desensitizes nicotinic acetylcholine receptors (nAChRs) in mesocorticolimbic brain circuitry involved in motivated behavior, volitional movement control, and reward prediction. Acute exposure of ventral tegmental area (VTA) neurons to nicotine depolarizes the membrane potential, enhances firing, and boosts dopamine release in target structures such as nucleus accumbens (NAc) (Brazell et al., 1990; Calabresi et al., 1989; Damsma et al., 1988; Grenhoff et al., 1986). Repeated exposure to nicotine induces critical neuroadaptations in these circuits (Mao et al., 2011; Saal et al., 2003), which underlie continued drug-seeking behavior and drug self-administration. Further elucidation of nicotine’s action in circuitry relevant to addiction is thus a high priority, as it could give rise to novel treatment strategies.

VTA DAergic and GABAergic neurons express β2-containing nAChRs in their somatodendritic compartments and presynaptic terminals in dorsal and ventral striatum (Giorguieff et al., 1977). Nicotine is understood to act on VTA circuitry by 1) directly stimulating DA neurons via nAChR activation (Calabresi et al., 1989), 2) inhibiting such neurons and other target brain structures through nAChR activation on VTA GABA neurons (Mansvelder et al., 2002), and 3) facilitating glutamate release from afferents endowed with homomeric α7 nAChRs (Mansvelder and McGehee, 2000). This framework has been valuable, but recent appreciation for VTA glutamate neurons, as well as DA neurons that co-release other transmitters (see (Morales and Margolis, 2017)), indicate that it is incomplete.

VTA DA neurons, defined either by DA transporter (DAT) or tyrosine hydroxylase (TH) expression, are preferentially found in lateral VTA subnuclei, make classical projections to target structures such as nucleus accumbens lateral shell and medial prefrontal cortex, and release DA in tonic and phasic patterns (Lammel et al., 2008; Tsai et al., 2009). GABAergic VTA neurons are defined by expression of glutamate decarboxylase 1 or 2 (GAD1/2). These cells are numerous in midline VTA (mVTA) subnuclei and project to a partially-overlapping set of target structures (Brown et al., 2012; Root et al., 2014b; Taylor et al., 2014) relative to DA neurons. VTA glutamate neurons have been examined directly via molecular targeting of vesicular glutamate transporter 2 (VGLUT2) (Tong et al., 2007; Vong et al., 2011). VGLUT2+ VTA neurons are, like GAD2(+) neurons, located preferentially in mVTA subnuclei (Yamaguchi et al., 2007) and project to ventral pallidum, lateral habenula, hippocampus, and nucleus accumbens medial shell (Hnasko et al., 2012; Ntamati and Luscher, 2016; Qi et al., 2016; Root et al., 2014a; Root et al., 2014b; Yoo et al., 2016). These circuits are functionally significant, being involved in both reward and aversion (Qi et al., 2016; Root et al., 2014a; Yamaguchi et al., 2007; Yoo et al., 2016). VTA VGLUT2+ neurons also make widespread intrinsic contacts with DAergic and non-DAergic neurons within the VTA itself (Dobi et al., 2010; Wang et al., 2015; Yoo et al., 2016).

Glutamatergic transmission in the VTA plays a critical role in nicotine dependence (Kenny et al., 2009), but the receptor and circuitry mechanisms are unclear. Whether and how nicotine modulates VTA output by impinging on intrinsic glutamatergic circuitry in VTA is completely unknown. Based on the influence of these neurons on local VTA circuitry, their distinct connectivity pattern with target structures, and their ability to co-release DA or GABA (Stamatakis et al., 2013; Stuber et al., 2010; Tecuapetla et al., 2010), we speculated that if nAChRs are functionally expressed in VGLUT2+ neurons, they could be uniquely positioned to influence motivated behavior circuits by enhancing excitatory transmission within, and downstream from, the VTA. In this study, we demonstrate the existence of functional nAChRs in VGLUT2+ neurons, characterize their pharmacological properties, examine their subcellular distribution pattern, and uncover a role for them in intrinsic excitatory transmission in VTA microcircuits.

## Results

### nAChRs are Expressed in Midline VTA neurons

We began investigating whether mVTA neurons participate in cholinergic transmission by asking whether they are innervated by cholinergic fibers. Mice expressing tdTomato (tdT) in cholinergic somata and axons were produced by crossing ChAT-IRES-Cre (ChAT-Cre) mice with Rosa26-LSL-tdTomato (Ai14) mice. mVTA interfascicular nucleus (IF) neurons in slices from ChAT-Cre::Ai14 mice were patch clamped, filled with Alexa 488, and imaged *in situ* via 2-photon laser scanning microscopy (2PLSM) (**Fig. 1a**). Proximal cholinergic fibers were also imaged in this regime. mVTA IF neurons were typically found adjacent to tdT+ fibers (**Fig. 1b**). Analysis of image stacks in xy, yz, and xz planes revealed a modest number of cholinergic fibers in close apposition to IF neuronal somata and dendritic processes (**Fig. 1c,d**), suggesting that these neurons are cholinoceptive. To examine this possibility, ChAT-Cre mice were microinjected in the pedunculopontine tegmental nucleus (PPTg) with Cre-dependent AAV-DIO-ChR2-EYFP vectors to enable photostimulation and release of ACh from cholinergic terminals in mVTA (**Fig. 1e**). We first verified that ChR2-EYFP was expressed selectively in ChAT+ PPTg neurons (**Fig. S1a**). We recorded robust 470 nm-stimulated (0.12 mW/mm^2^) photocurrents in voltage clamp recordings from ChR2-EYFP+ PPTg neurons using a variety of stimulation parameters (**Fig. S1b-d**). In current clamp recordings, photostimulation (0.1 ms) triggered single action potentials (**Fig. S1e**). ChR2-EYFP+ cholinergic fibers were evident in close proximity to mVTA IF neurons (**Fig. S1f**), but were not found in uninfected mice. To examine cholinergic transmission in mVTA through nAChRs, voltage-clamped IF neurons in slices containing ChR2-EYFP+ fibers were photostimulated with 470 nm light flashes (20 s, 0.12 mW/mm^2^). In this regime, a slow inward current was activated (**Fig. 1f,g**), suggestive of nAChR activity (Xiao et al., 2016). If these currents are nAChR-dependent, they should be augmented by galantamine (nAChR positive allosteric modulator) and inhibited by nAChR antagonists. After validating galantamine’s ability to enhance slow ACh-stimulated currents in IF neurons (**Fig. S1g,h**), we recorded 470 nm-stimulated photocurrents before and after galantamine (1 μM) bath application. Galantamine significantly enhanced the amplitude of these photocurrents (**Fig. 1h,i**). Subsequent co-application of a nAChR antagonist cocktail with galantamine eliminated these photocurrents (**Fig. 1i**). Collectively, these 2PLSM and optogenetic studies indicate that mVTA IF neurons are innervated by cholinergic fibers and participate in cholinergic transmission through nAChRs.

**Figure 1.**
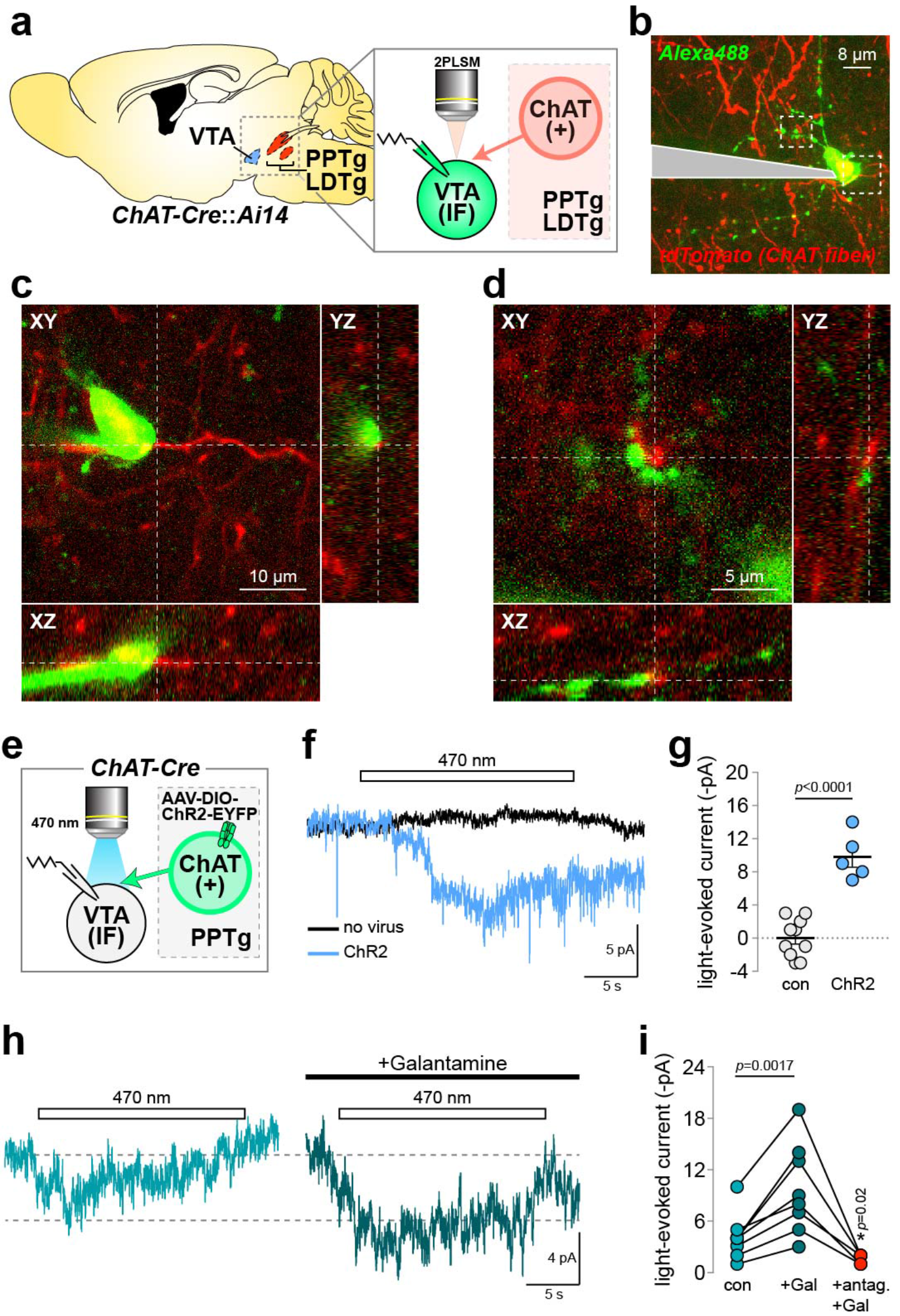
Midline VTA nAChRs participate in cholinergic transmission. **(a)** Schematic showing VTA and PPTg/laterodorsal tegmental nucleus (LDTg). Inset: ChAT-Cre::Ai14 mice studied in (b-d). mVTA neurons and tdT+ cholinergic fibers were imaged via 2PLSM during patch clamp recordings. **(b)** Maximum intensity projection of a z-series 2PLSM image for a mVTA neuron filled with Alexa488 (green channel) during patch clamp recording. Local cholinergic fibers are shown in the red channel. **(c,d)** The soma (c) and dendrites (d) from the cell shown in (b) are shown at higher magnification. XY/XZ/YZ planes from z-series 2PLSM merged (Alexa488and tdT channels) image show cholinergic fibers in close proximity to the mVTA neuron soma (c) and dendrites (d). Representative of n=4/3. **(e)** Optical approach used in (f-i). ChAT-Cre mice were microinjected in PPTg with AAV expressing Cre-dependent ChR2, patch clamp recordings were made in mVTA neurons, and ChR2 activity was controlled via epi-illumination flashes (470 nm, 0.12 mW/mm^2^) through the microscope objective. **(f-g)** Slow cholinergic photocurrents in mVTA neurons. (f) Voltage clamp recordings showing current deflections during 470 nm light flashes (20 s) in slices from ChR2- vs. ChR2+ mice. Averaged traces for 5-10 independent cells are shown. (g) Light-evoked current amplitudes are plotted for (n=9/2 and 5/2) individual neurons in control (no virus) and ChR2+ mice. *P* value: unpaired *t*-test. **(h,i)** Pharmacological analysis of cholinergic photocurrents. (h) Averaged photocurrents (470 nm, 20 s flash) are shown for neurons from ChR2+ mice before (left trace) and after (right trace) bath application of galantamine (1 μM). (i) Light-evoked current amplitudes are plotted for (n=8/4) individual neurons before drug (con), during galantamine (Gal) application, and during co-application of galantamine and a nAChR antagonist cocktail containing (10 μM DHβE, 100 nM α-Ctx MII, 10 nM MLA). *P* values: paired *t*-tests. See also **Fig. S1**, **S2**.

To examine nAChR subunit expression in mVTA and corroborate these physiology results, mice expressing GFP-tagged nAChR subunits were employed (Shih et al., 2014). Anti-GFP immunostaining suggested expression of α4, α6, β2, and β3 nAChR subunits in mVTA (**Fig. S2a**). There was little to no evidence for α3 and β4 subunit expression therein (**Fig. S2a**). α6 expression was strongest in mVTA rostral linear (RLi) and IF nuclei, and many α6+ neurons were also positive for TH (**Fig. S2b**). Additionally, there were several α6+ neurons that lacked TH as well as TH+ neurons lacking α6 (**Fig. S2b**). These data suggest that several nAChR subunits are expressed in mVTA nuclei known to be enriched in VGLUT2+ and GAD2+ neurons.

### mVTA glutamate neurons express functional nAChRs

To directly examine nAChR subunit expression in VTA glutamate neurons, triple-channel fluorescence mRNA *in situ* hybridization (FISH) was used. First, nAChR probes for *Chrna4, Chrna6*, and *Chrnb2* were validated by comparing lateral VTA *Th* (tyrosine hydroxylase) colabeling patterns with a previously published *in situ* hybridization study (Azam et al., 2002) (**Fig. S3**). Co-labeling of mVTA IF neurons with probes for *Slc17a6* (VGLUT2), *Chrna4*, and *Th* revealed neurons with multiple co-expression patterns (**Fig. 2a**). Quantification of these results indicated that a large majority of *Slc17a6+* neurons are positive for *Chrna4* (**Supplementary Table 1; Fig. 2b,c**). Examining *Th* expression within *Slc17a6*+/*Chrna4*+ neurons indicated that a majority of these neurons also express *Th* (**Supplementary Table 1**; **Fig. 2c**). For comparison, the same analysis was performed on lateral VTA neurons in the parabrachial pigmented nucleus (PBP). Co-labeling of *Slc17a6* and *Chrna4* was qualitatively similar in these neurons, as was the fraction of *Slc17a6*+/*Chrna4*+ neurons that co-labeled for *Th* (**Supplementary Table 1**; **Fig. 2d,e**). The same analysis was performed using probes for *Chrna6* and *Chrnb2*, and produced very similar results (**Fig. 2f-o**). To determine whether the nAChR expression profile in VTA glutamate neurons is unique or generalizable, FISH was performed examining nAChR expression in GABA (*Gad2*+) neurons (**Fig. S4a,f,k**). Like *Slc17a6* labeling revealed, the vast majority of IF *Gad2*+ cells also express *Chrna4, Chrna6*, and *Chrnb2* (**Fig. S4b,c,g,h,l,m**). Surprisingly, *Th* was expressed in 30-50% of Gad2+/nAChR+ neurons, depending on the subunit examined (**Fig. S4c,h,m**). Compared to mVTA IF, lateral VTA PBP neurons tended to have slightly fewer neurons co-labeling for *Gad2* and nAChR subunits as well as fewer *Th*+ neurons that co-labeled for *Gad2* and nAChR subunits (**Fig. S4d,e,i,j,n,o**).

**Figure 2.**
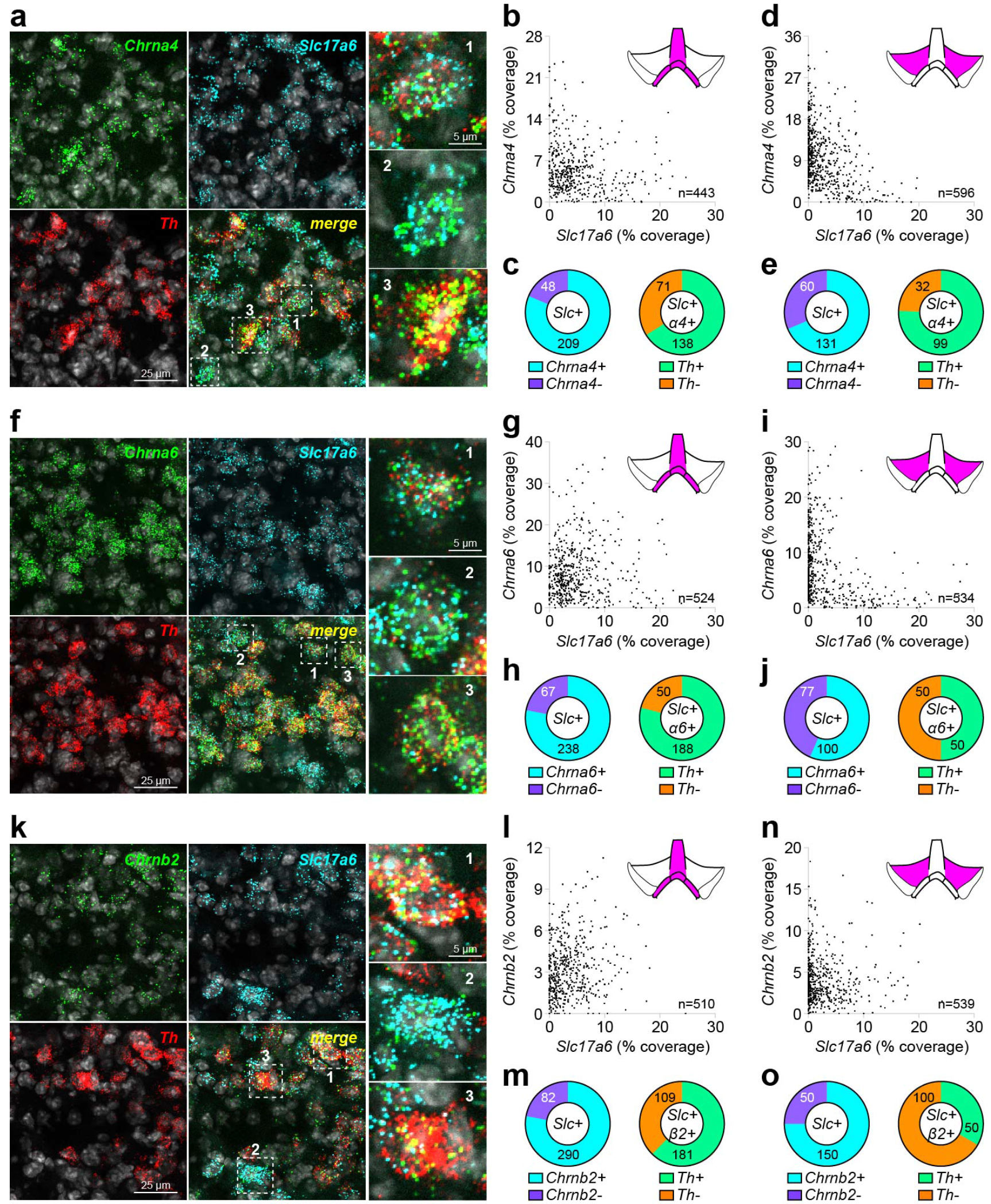
VTA glutamate neurons express heteromeric nAChRs. **(a-e)** Co-expression of *Chrna4* and *Slc17a6* in VTA neurons. **(a)** Representative 3-channel fluorescence *in situ* hybridization (FISH) images in mVTA neurons for probes: *Chrna4*, *Slc17a6* (VGLUT2), and *Th*. Single *Chrna4*+ neurons indicated in the ‘merge’ panel, and shown enlarged (right panels), had this expression profile: 1 - *Chrna4*+/*Slc17a6*+/*Th*+; 2 - *Chrna4*+/*Slc17a6*+/*Th*-; 3 - *Chrna4*+/*Slc17a6*-/*Th*+. **(b-c)** Analysis of *Chrna4/Slc17a6* co-expression in mVTA. **(b)** Scatter plots showing *Slc17a6* (abscissa) and *Chrna4* (ordinate) coverage’ for all analyzed neurons. **(c)** Left: Pie graph showing the fraction of *Slc17a6*+ neurons that were *Chrna4*+ vs. *Chrna4*-. Right: *Th* expression profile is shown for *Slc17a6*+/*Chrna4*+ neurons from left pie graph. **(d-e)** Analysis of *Chrna4*/*Slc17a6* co-expression in lateral VTA was performed identically as in mVTA (b-c). **(f-j)** Co-expression of *Chrna6* and *Slc17a6* in VTA neurons. Micrographs and analysis for *Chrna6* were performed as for *Chrna4*. **(k-o)** Co-expression of *Chrnb2* and *Slc17a6* in VTA neurons. Micrographs and analysis for *Chrnb2* were performed as for *Chrna4* and *Chrna6*. Complete data for nAChR/*Slc17a6* FISH is listed in **Supplementary Table 1**. Data are pooled from n=3 mice. See also **Fig. S3**, **S4** and **Supplementary Table 2**.

To directly probe functional nAChRs in VTA glutamate neurons in IF, we produced mice expressing tdT in VGLUT2+ neurons (VGLUT2-Cre::Ai14 mice) to enable targeted recordings in slices. In parallel, we created mice enabling targeted recordings in VTA DA and GABA neurons using DAT-Cre and GAD2-Cre mice crossed with Ai14 mice. VTA sections from this trio were costained with anti-DsRed and anti-TH antibodies to examine/validate the tdT expression profile (**Fig. 3a,d,g**). In voltage-clamp, we recorded robust ACh-elicited currents in VGLUT2+ IF neurons (Fig. 3b). Using stepwise application of pharmacological agents in conjunction with ACh pressure ejection application to examine the nAChR subtypes, we found evidence for α6* (*=containing the indicated subunit and possibly others) and α4β2(non-α6) nAChRs (**Fig. 3c**). In pooled C57Bl/6 but not VGLUT2+, GAD2+, or DAT+ VTA neurons (paired *t*-tests, respectively: *p*=0.0283, *p*=0.1136, *p*=0.0860, *p*=0.1327), an α7 nAChR antagonist (methyllycaconitine (MLA); 10 nM) further reduced the residual ACh-evoked current that was insensitive to combined α-Ctx MII[H9A;L15A]+DHβE treatment. A cocktail containing 10 μM CNQX, 50 μM AP5, 100 μM picrotoxin, and 0.5 μM TTX partially reduced the ACh-elicited currents in VTA VGLUT2+ neurons, suggesting a minor contribution from presynaptic nAChRs on glutamatergic terminals (**Fig. 3j**). A control experiment demonstrated that run-down of nAChR currents cannot account for the reduction in current we observed with long (60 min) recordings during pharmacological analysis (**Fig. 3k**). We examined ACh-elicited currents in VTA GABA (**Fig. 3d-f**) and DA (**Fig. 3g-i**) neurons in an identical manner and found qualitatively similar results. To control for any potential strain differences in nAChR function, we performed a similar analysis in unlabeled VTA IF neurons from C57BL/6 mice. Again, these results from a presumptive pool of DA/GABA/VGLUT2 neurons were qualitatively similar to results in the Cre lines (**Fig. 3l,m**).

**Figure 3.**
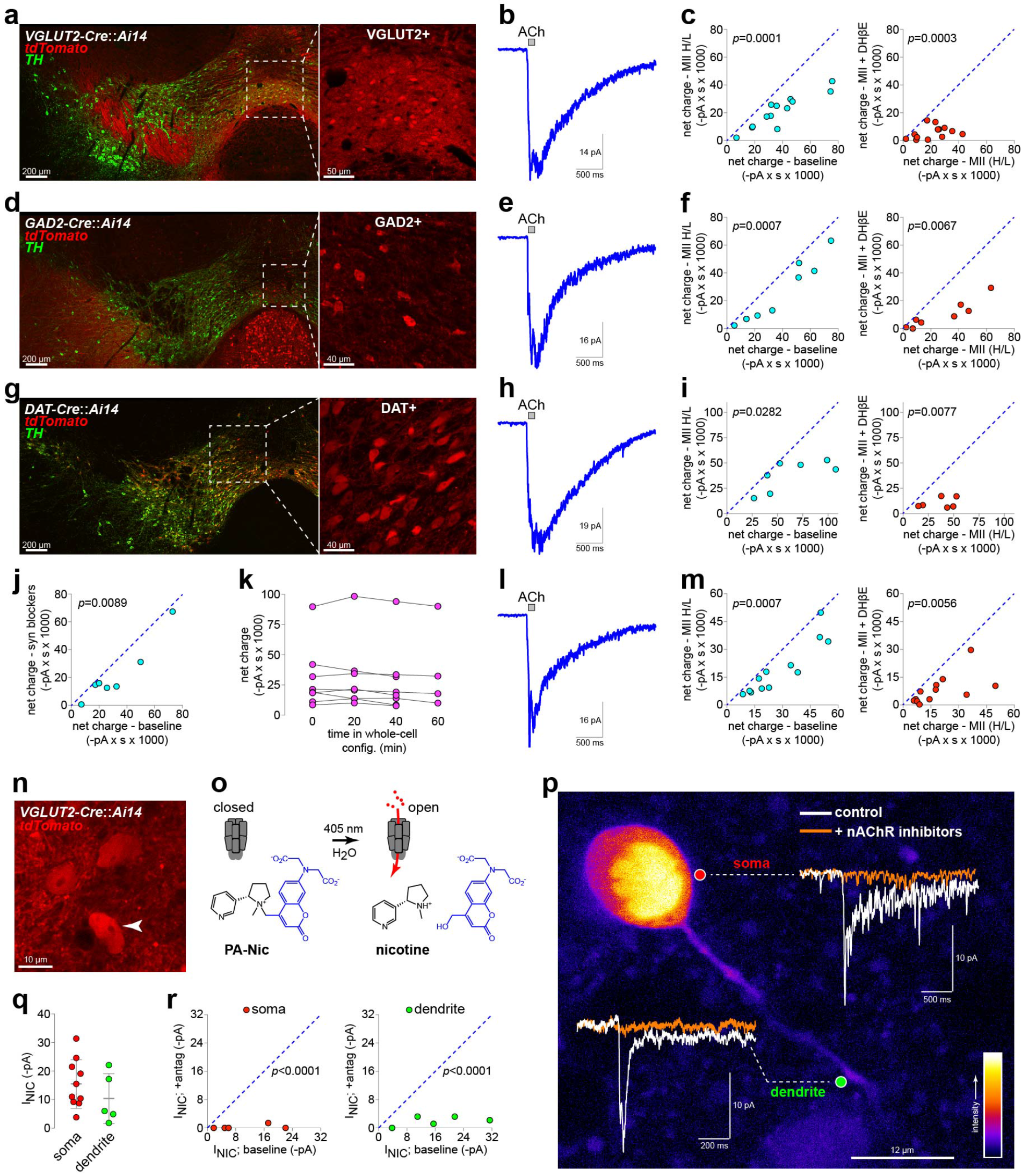
Functional nAChRs in VTA glutamate neurons. **(a,d,g)** Reporter mouse strains for targeted recordings in neurotransmitter-defined VTA cell types. Rosa26-LSL-tdTomato (A114) mice were crossed to VGLUT2-Cre (a), GAD2-Cre (d), and DAT-Cre (g), and images are shown of coronal VTA sections from progeny mice stained with anti-TH and anti-DsRed antibodies. Right panel: tdTomato-only channel of boxed area showing VGLUT2+ (a), GAD2+ (d), and DAT+ (g) neurons. **(b,e,h,l)** Averaged trace is shown for pressure ejection applications of 1 mM ACh to voltage clamped VTA VGLUT2+ (b), GAD2+ (e), DAT+ (h), and mixed mVTA neurons in C57BV6 mice (l). **(c,f,I,m)** Left: plot of 1 mM ACh-induced net chargeat baseline vs. after bath application of α-Ctx MII[H9A;L15A] for n=8/3 individual VGLUT2+ (c), n=7/2 GAD2+ (f), n=5/2 DAT+ (i), and n=9/5 mixed C57Bl/6 (m) neurons. Right: plot of 1 mM ACh-induced net charge in the presence of α-Ctx MII[H9A;L15A] vs. after bath application of α-Ctx MII[H9A;L15A] + 1 μM DHβE for the same neurons shown at left. *P* values: paired *t*-test. **(j)** Presynaptic contribution to ACh-evoked currents. A plot of 1 mM ACh-induced net charge at baseline vs. after bath application of a cocktail containing CNQX (10 μM), D-AP5 (50 μM), TTX (0.5 μM), and picrotoxin (100 μM). n=7/2. *P* value: paired *t*-test. **(k)** ACh-evoked currents rundown analysis. ACh net charge responses to repeated pressure ejection application of 1 mM ACh is shown for n=8/2 individual mVTA VGLUT2+ neurons. Responses were recorded at the indicated times after establishing a whole-cell recording. **(n)** VGLUT2(+) neurons (solid arrowhead) in VGLUT2-Cre::Ai14 mice were imaged via 2PLSM and nicotine photolysis was executed during patch clamp recordings. **(o)** Nicotine uncaging schematic. A photoactivatable nicotine probe (PA-Nic) was applied to mVTA glutamate neurons, and nicotine was uncaged with focal 405 nm laser photolysis. **(p)** 2PLSM image of a patch clamped mVTA glutamate neuron is shown, along with overlaid example current traces of nicotine-evoked current responses following nicotine uncaging. Somatic and dendritic responses were evoked before (white trace) and after (orange trace) bath application of a nAChR antagonist cocktail (10 μM DHβE, 100 nM α-Ctx MII, 100 nM MLA). **(q,r)** Analysis of nicotine uncaging responses. (q) Response amplitudes are shown, along with mean and standard deviation, for nicotine uncaging at the soma (n=10/7) and dendrite (n=5/4) of individual glutamate neurons. (r) Somatic and dendritic nicotine-evoked currents are blocked by bath application of a nAChR antagonist cocktail (same as in (p)). *P* value: paired *t*-test on normalized data.

Next, we asked whether nAChRs in mVTA glutamate neurons are localized to somata, dendrites, or both. VGLUT2+ IF neurons in slices from VGLUT2-Cre::Ai14 mice were identified via tdT fluorescence using 2PLSM imaging (**Fig. 3n**). A photoactivatable nicotine (PA-Nic) probe (REFERENCE), which releases nicotine following 405 nm laser flash photolysis (**Fig. 3o**), was used to study functional nAChR responses in mVTA VGLUT2+ neurons during recording and 2PLSM imaging. PA-Nic photolysis in spots (1 μm diameter) immediately adjacent to VGLUT2+ neuron somata and dendrites revealed fast inward currents of modest amplitude (**Fig. 3p,q**). These currents were mediated by nAChRs, as an antagonist cocktail (10 μM DHβE, 100 nM α-Ctx MII, 100 nM MLA) eliminated these responses (**Fig. 3r**). These data provide definitive proof for the existence of functional somatodendritic nAChRs in mVTA glutamate neurons.

### mVTA glutamate neuron nAChRs modulate excitatory neurotransmission

In addition to their long-range projections, VTA glutamate neurons make excitatory connections with local VGLUT2-neurons in VTA (Wang et al., 2015; Yoo et al., 2016). We asked whether nAChR activity in VGLUT2+ neurons can modulate glutamatergic transmission in this microcircuit. ChR2 was selectively expressed in mVTA VGLUT2+ neurons by microinjecting Cre-dependent AAV-DIO-ChR2-EYFP (or mCherry) vectors in mVTA of VGLUT2-Cre mice. Expression of ChR2-EYFP in VGLUT2+ neurons was first validated by injection of AAV-DIO-ChR2-EYFP into mVTA of VGLUT2-Cre::Ai14 mice with tdT-marked VGLUT2+ neurons and fibers. Immunohistochemistry analysis revealed that our microinjections saturated the mVTA with ChR2+ fibers and infected VGLUT2+ cell bodies (**Fig. 4a**); all tdT+ neurons in mVTA of n=3 mice co-expressed ChR2-EYFP. Based on this analysis, we conclude with a high degree of certainty that ChR2-lacking neurons studied below are not VGLUT2+ neurons.

**Figure 4.**
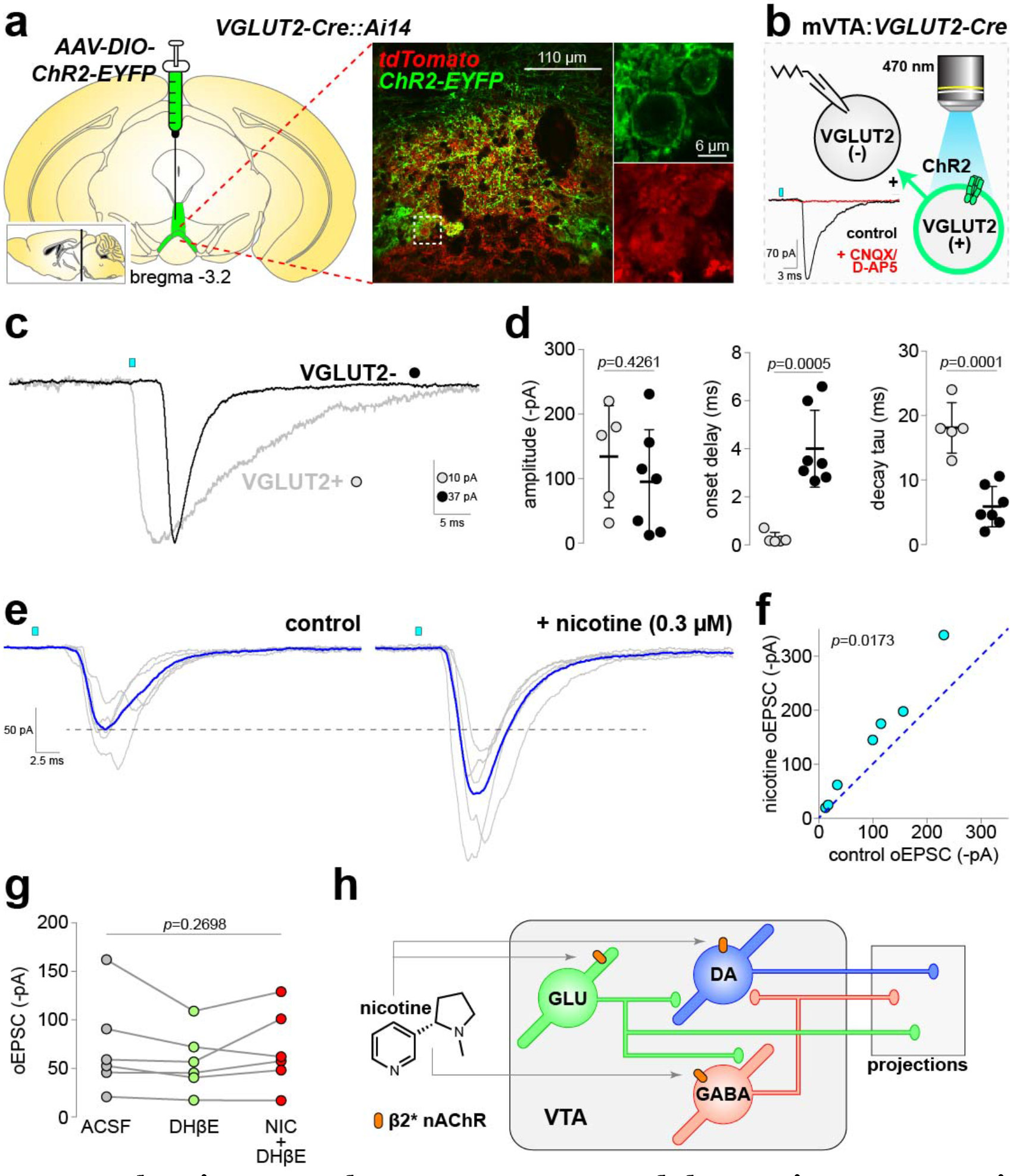
nAChRs in mVTA glutamate neurons modulate excitatory transmission. **(a)** Validation of ChR2 expression in mVTA glutamate neurons. AAVs directing Cre-dependent ChR2-EYFP (or mCherry) expression were unilaterally microinjected into mVTA of VGLUT2-Cre::Ai14 mice, and sections were double-stained with anti-DsRed and anti-GFP antibodies to observe VGLUT2+ neurons expressing ChR2 (large image). Enlarged views (right) of the boxed area show VGLUT2+ neuron expression of ChR2-EYFP. **(b)** Optically-evoked synaptic currents in mVTA. In mVTA slices from VGLUT2-Cre mice microinjected as described in (a), uninfected VGLUT2-neurons were patch-clamped and optical EPSCs were recorded. oEPSCs were sensitive to CNQX+AP5 (lower left). **(c-d)** Photocurrents in VGLUT2+ neurons and oEPSCs in VGLUT2- neurons are distinct. A representative oEPSC and photocurrent in VGLUT2- and VGLUT2+ neuron, respectively, are shown (c). Analysis of photocurrent and oEPSC amplitude, onset delay, and decay tau in n=5/4 VGLUT2+ and n=7/5 VGLUT2-mVTA neurons (d). *P* values: unpaired *t*-test. **(e-f)** Nicotine enhances VGLUT2-dependent oEPSCs in mVTA. Representative oEPSC recordings (individual trials=grey; averaged trace=blue) are shown for VGLUT2- neurons before (control) and after nicotine (0.3 μM) bath application (e). **(f)** Summary plot of control vs. nicotine-enhanced oEPSC peak current amplitude (n=7/5). *P* values: paired *t*-test. **(g)** Before-after plot showing oEPSC amplitude in VGLUT2- neurons (n=6/6) during bath application of ACSF (control), ACSF + DHβE (1 μM), and ACSF + DHβE (1 μM) + nicotine (0.3 μM). *P* value: repeated measures 1-way ANOVA. **(h)** Model for nicotine action at somato-dendritic nAChRs in VTA. See also **Fig. S5**.

Next, we recorded from local uninfected neurons in mVTA to study excitatory transmission between VGLUT2- and ChR2-expressing VGLUT2+ neurons (**Fig. 4b**). Full field illumination (470 nm, 1-5 ms) synchronously activated ChR2 on VGLUT2+ somata and local terminals, evoking optical EPSCs (oEPSCs) in VGLUT2- mVTA neurons (**Fig. 4b**, **lower left**). oEPSCs were sensitive to bath-applied CNQX+AP5 (**Fig. 4b**), indicating they are synaptic in nature. Some VGLUT2+ neurons expressed weak levels of ChR2-EYFP in their somata, but oEPSCs in VGLUT2-neurons were distinguishable from direct photocurrents in VGLUT2+ neurons (**Fig. 4c**) via their clear synaptic delay and faster decay time constant (**Fig. 4d**). To examine whether nAChRs in VGLUT2+ mVTA neurons modulate these oEPSCs, we compared the oEPSC amplitude before and after bath application of nicotine (0.3 μM). Optical pulse duration (0.5-5 ms) and flash strength (<0.01 mW/mm^2^) were empirically chosen for each cell such that baseline responses were initially ~50-150 pA (**Fig. 4d,e**). Nicotine application significantly enhanced oEPSC amplitude in mVTA VGLUT2- neurons (**Fig. 4e,f**), demonstrating a role for nAChRs in excitatory transmission between mVTA VGLUT2+ and VGLUT2− neurons. Nicotine-modulated oEPSCs may occur in mVTA DAT+ and/or GAD2+ neurons, but these neuron types were not uniquely identifiable when examining I_*h*_ current amplitude, action potential firing rate, or input resistance (**Fig. S5**). Finally, nicotine (0.3 μM) was unable to enhance oEPSC amplitudes in the presence of DHβE (RM 1-way ANOVA of ACSF vs. ACSF + DHβE vs. ACSF + DHβE + nicotine; *F*(1.577,7.885)=1.523, *p*=0.2698) (**Fig. 4g**), implicating β2* heteromeric nAChRs in mVTA glutamate neuron excitatory transmission.

## Discussion

We have used a diverse set of molecular, genetic, and physiological approaches to make the following observations regarding VTA glutamate neuron nAChRs. First, most VTA glutamate neurons co-express heteromeric nAChRs, including many neurons that are also TH+. These receptors are functional and largely indistinguishable from nAChRs on DAT+ and GAD2+ VTA cells. Second, optical release of ACh from cholinergic fibers demonstrates that these receptors are functionally integrated into cholinergic circuits. Nicotine photolysis demonstrates nAChR functionality on somata and dendrites, and additional optogenetic experiments demonstrate that these receptors modulate excitatory transmission in VTA microcircuits.

In this study, we studied how nAChR distribution and function maps onto the recently-appreciated trio (DAT, GAD2, VGLUT2) of neurotransmitter-defined VTA neurons by examining these receptors in glutamate neurons and comparing their expression and function to nAChRs in DA and GABA neurons. Heteromeric β2* nAChRs, which exhibit high sensitivity to ligand (Salminen et al., 2007) and strong nicotine-mediated desensitization (Pidoplichko et al., 1997), are the predominant subtype in these cells. This is consistent with prior studies of nAChRs in putative DA neurons (Azam et al., 2002; Klink et al., 2001), but extends those findings by suggesting not only nicotinic cholinergic modulation of glutamate release from VTA neurons, but also similar modulation of DA/glutamate (Stuber et al., 2010) and GABA/glutamate (Root et al., 2014b; Yoo et al., 2016) co-release. Given that VTA VGLUT2+ neurons project to atypical forebrain targets, cholinergic modulation of VTA-derived glutamatergic afferents may occur at synapses that are not yet characterized. Consistent with the relatively non-specific targeting of VTA DAT/GAD2/VGLUT2 neurons by hindbrain cholinergic nuclei (Faget et al., 2016), we did not find evidence for selective expression of any particular nAChR subtype in these neuron types. The caveat to this is that while the group average did not indicate specificity of nAChR subtype expression, individual VGLUT2+, DAT+, and GAD2+ cells express differing levels of functional nAChR subtypes (**Fig. 3**).

Our results showing that midline VTA IF neurons are innervated by cholinergic fibers extends prior work in ChAT-Cre rats that did not identify which VTA subnuclei are activated by cholinergic afferents (Dautan et al., 2016; Xiao et al., 2016). Results from 2PLSM imaging (**Fig. 1b-d**), and confirmed in mVTA sections containing ChR2-EYFP ChAT+ fibers (Fig. S1a), show modest to sparse innervation of individual IF neurons by cholinergic fibers. This is generally consistent with the slow kinetics of inward currents activated by optical release of ACh. Prolonged photostimulations (20 s) were required to elicit these currents, and we found no evidence for fast (ms time scale) cholinergic transmission evident in other brain areas (Estakhr et al., 2017; Hedrick and Waters, 2015). Based on this, volume transmission may be the operable cholinergic mechanism in mVTA. Despite this likelihood, we also uncovered fast (ms timescale) nAChR currents at VTA VGLUT2+ neuronal somata and dendrites when rapidly “jumping” the concentration of exogenous nicotine using laser photolysis of PA-Nic (**Fig. 3p-r**). These experiments strongly suggest that the slow responses following optical release of ACh are related to the distance ACh travels – while evading metabolism by acetylcholinesterase – between its release site and its target receptor. Our nicotine photolysis results also confirm what has previously only been inferred: functional VTA neuron nAChRs are not only located on the soma where they likely play a role in modulating action potential firing (Calabresi et al., 1989), but also in dendrites where they may mediate cholinergic modulation of dendritic integration (see (Oldenburg and Ding, 2011)). Lastly, these nicotine photolysis studies, juxtaposed with optical release of ACh, illustrate the dramatic difference between endogenous ACh transmission and how it is biophysically “highjacked” by nicotine.

VTA GABA neurons have long been known to send local collaterals to neighboring DA cells (Johnson and North, 1992), but more recently a local glutamatergic connection between VGLUT2+ and TH+ neurons has been described that modulates reward processing (Wang et al., 2015; Yoo et al., 2016). The present work extends those results by revealing that nicotine can potently enhance excitatory transmission at these synapses (**Fig. 4h**). In doing so, nicotine increases the gain and reliability of local glutamate to DA neuron transmission in VTA. Additionally, by this activity nicotine will likely sensitize mVTA DA and/or GABA neurons to other excitatory input, such as corticolimbic glutamate transmission (Mansvelder and McGehee, 2000) and direct somatodendritic nAChR stimulation by nicotine or ACh. Nicotine’s reinforcing property, along with its ability to modulate stress responses (Morel et al., 2017), is critically dependent on its interaction with VTA circuitry (Corrigall et al., 1994). Until now, the totality of nicotine’s action in VTA was assumed to involve activation of nAChRs on DA and GABA neurons coupled with potentiation of extra-VTA glutamatergic and GABAergic inputs (Champtiaux et al., 2002; Klink et al., 2001; Mansvelder et al., 2002; Mansvelder and McGehee, 2000). Our demonstration that nAChRs on VTA glutamate neurons are involved in nicotine modulation of local microcircuit activity adds an additional level of complexity to our understanding of nicotine’s pharmacological action in mesolimbic circuits. These results require that we update the existing framework regarding nicotine-elicited modulation of electrical and chemical signaling in the mesocorticolimbic system (**Fig. 4h**).

## Acknowledgements

This work was supported by National Institutes of Health (NIH) grants (DA035942, DA040626, and DA030396 to R.M.D.; GM103801 and GM48677 to J.M.M.; DA028955 to Henry A. Lester), the Howard Hughes Medical Institute, and funds from Northwestern University. M.C.A. was supported by a fellowship from the PhRMA Foundation. D.L.W. was supported by the JPB Foundation. Thanks to the following principle investigators and their laboratories at Northwestern University for excellent technical assistance, instrumentation help, manuscript discussion, and mouse strains: D. James Surmeier, Savio Chan, Loukia Parisiadou, Yevgenia Kozorovitskiy, Anis Contractor. Thanks to the lab of Henry A. Lester (California Institute of Technology) for providing mouse strains.

## Author Contributions

Conceptualization, R.M.D.; Methodology, C.P., M.C.A., X-T.J., Y.W., D.L.W., and R.M.D.; Software, R.M.D.; Validation, Y.Y., C.P., M.C.A., X-T.J., M.D.R., Y.W., and D.L.W.; Formal Analysis, R.M.D.; Investigation, Y.Y., C.P., M.C.A., X-T.J., Y.W., and R.M.D.; Resources, S.B., V.J.K., L.D.L., and J.M.M.; Data Curation, R.M.D.; Writing – Original Draft, R.M.D.; Writing – Review & Editing, Y.Y., C.P., M.C.A., X-T.J., Y.W., D.L.W., J.M.M., and R.M.D.; Visualization, R.M.D.; Supervision, R.M.D.; Project Administration, R.M.D.; Funding Acquisition, R.M.D.

## Materials and Methods

### Materials and Viral Vectors

AAV9.EF1a.DIO.hChR2(H134R)-eYFP.WPRE.hGH and AAV9.EF1a.DIO.hChR2(H134R)-mCherry.WPRE.hGH were obtained from Penn Vector Core. α-conotoxins were synthesized as previously described (Azam et al., 2010). Dihydro-β-erythroidine hydrobromide (DHβE), methyllycaconitine (MLA), picrotoxin, and atropine sulfate (atropine) were obtained from Sigma. 6-Cyano-7-nitroquinoxaline-2,3-dione (CNQX), D-(-)-2-Amino-5-phosphonopentanoic acid (D-AP5), Octahydro-12-(hydroxymethyl)-2-imino-5,9:7,10a-dimethano-10aH-[1,3]dioxocino[6,5-d]pyrimidine-4,7,10,11,12-pentol (TTX), QX314 chloride (QX314), and galantamine hydrobromide (galantamine) were obtained from Tocris. Alexafluor (Alexa) 488 was obtained from Life Technologies. PA-Nic was synthesized as previously described (REFERENCE).

### Mice

All experimental protocols involving mice were reviewed and approved by the Institutional Animal Care and Use Committee at Northwestern University (protocols #IS00003282, IS00003604). Procedures also followed the guidelines for the care and use of animals provided by the National Institutes of Health Office of Laboratory Animal Welfare. All efforts were made to minimize animal distress and suffering during experimental procedures, including during the use of anesthesia. Mice were housed at 22°C on a 12-hour light/dark cycle with food and water *ad libitum*. Mice were weaned on postnatal day 21 and housed with same-sex littermates. A tail sample was taken from each mouse for genotyping via polymerase chain reaction (PCR) as previously described (Drenan et al., 2008; McGranahan et al., 2011). Creation and characterization of knock-in (α3-GFP, α4-GFP, β2-GFP, β3-GFP, β4-GFP) or bacterial artificial chromosome transgenic mice (α6-GFP) harboring in-frame insertions of GFP into the coding sequence of specific nAChR subunit genes have been previously described (Mackey et al., 2012; Shih et al., 2014). The following mouse strains were obtained from Jackson Laboratories: ChAT-IRES-Cre (Jax #006410) (Rossi et al., 2011), VGLUT2-IRES-Cre (Jax #016963) (Vong et al., 2011), DAT-IRES-Cre (Jax #06660) (Backman et al., 2006), GAD2-IRES-Cre (Jax #010802) (Taniguchi et al., 2011), Ai14 (Jax #007908) (Madisen et al., 2010), C57Bl/6J (Jax #000664). Mice expressing tdT in a Cre-dependent manner (ChAT-IRES-Cre::Ai14, VGLUT2-IRES-Cre::Ai14, GAD2-IRES-Cre::Ai14, DAT-IRES-Cre::Ai14) were obtained by crossing mice heterozygous for each mutation, which produced ~25% double-heterozygous progeny. Male and female mice were used in approximately equal numbers.

### Stereotaxic Surgery

Male and female mice were used for surgery starting at 8 weeks of age. Mice were initially anesthetized with an intraperitoneal (i.p.) injection of a ketamine/xylazine mixture (120 mg/kg ketamine, 16 mg/kg xylazine). Mice were given additional “boost” injections of ketamine (100 mg/kg, i.p.) as needed. Alternatively, some mice were anesthetized with isoflurane: 3% (flow rate 500 mL/min) for induction and 1.5% (28 mL/min) for maintenance. Mice were secured into a stereotaxic frame and a small incision at the top of the head was made to expose the skull. Coordinates (unilateral) used for mVTA injections were (relative to bregma, in mm): M/L: +0.01 (or −0.01), A/P: −3.2, D/V: −4.55. Coordinates (bilateral) used for lateral VTA injections were (relative to bregma, in mm): M/L: ±0.5, A/P: −3.2, D/V: −4.75. Coordinates (bilateral) used for PPTg were (relative to bregma, in mm): M/L: ±1.20, A/P: −4.45, D/V: −4.05. Exact coordinates were adjusted to account for slight differences in the head size of individual mice: the bregma/lambda distance measured for each mouse was divided by the reported bregma/lambda distance for C57 mice (4.21), then multiplied by the A/P coordinate. The injection needle was slowly lowered through the drilled hole to the D/V coordinate. For AAV viruses, 500 nL of virus was infused at a rate of 50 nL/min. The injection needle was left in place for 10 min after the infusion ended before slowly retracting the needle. Sutures were used to close the incision. At the conclusion of the surgery, mice were given ketoprofen (5 mg/kg, s.c.) and placed in a recovery cage, kept warm, and observed until they were ambulatory. Mice were single-housed following virus injection surgery and were given at least 14 days to recover and for the virus to express before beginning experimental procedures. For electrophysiology experiments, accurate targeting of VTA was determined via direct visualization of fluorescent neurons in brain slices during recordings. Slices without such neurons were not used for recordings.

### Immunohistochemistry and Confocal Microscopy

Anti-GFP immunostaining and visualization with 3,3’-diaminobenzidine (DAB) of nAChR-GFP knockin/transgenic mouse brains (**Supplementary Fig. 2a**) was performed as part of the same tissue analysis as we previously described (Shih et al., 2014). All other immunohistochemistry in the paper was perfomed as follows. Mice were anesthetized with sodium pentobarbital (200 mg/kg, i.p.) and transcardially perfused with 10 mL of heparin-containing phosphate buffered saline (PBS) followed by 30 mL of 4% paraformaldehyde. Brains were dissected and postfixed in 4% paraformaldehyde overnight at 4°C. Coronal brain slices (50 μm) were cut on a freezing sliding microtome (SM2010R; Leica). VTA-containing slices were stained using the following procedure. Slices were first permeabilized for 2 min via incubation in PBST (0.3% Triton X-100 in PBS), followed by a 60 min incubation in blocking solution (0.1% Triton X-100, 5% horse serum in Tris-buffered saline (TBS)). Primary antibodies used in this study were as follows: sheep anti-TH (Millipore AB1542), rabbit anti-GFP (Invitrogen A11122), rabbit anti-DsRed (Clontech 632496), goat anti-ChAT (Millipore AB144P). Primary antibodies were diluted in blocking solution (anti-TH at 1:800, anti-GFP at 1:500, anti-DsRed at 1:500, anti-ChAT at 1:100). Slices were incubated in primary antibodies overnight at 4°C. Three 5 min washes in TBST (0.1% Triton X-100 in TBS) were done before transferring slices to secondary antibodies for a 60 min incubation at room temperature (anti-sheep or anti-rabbit Alexa 555, anti-rabbit Alexa 488, diluted to 1:500 in blocking solution). Slices were washed as before, mounted on slides, and coverslipped with Vectashield. Staining in the VTA was imaged as previously described (Mackey et al., 2012) with a Nikon A1 laser-scanning confocal microscope.

### mRNA fluorescence in situ hybridization and image analysis

Mice were deeply anesthetized with sodium pentobarbital (200 mg/kg, i.p.) and decapitated. Brains were quickly removed on ice, snap frozen, and embedded in cryo-embedding medium (OCT). Brains were sectioned on a cryostat (CM3050; Leica) into 20 μm sections, sections were adhered to Superfrost® Plus slides, and kept at −20°C to dry for 60 min and stored at −80°C until use. Sections were fixed with 4% paraformaldehyde and processed for RNAscope (Advanced Cell Diagnostics) multichannel fluorescent *in situ* hybridization (FISH) according to the manufacturer manual for Multiplex assays. Sections were counterstained with DAPI for 30 s at room temperature. Probes for detection of specific targets (*Chrna4*, *Chrna6*, *Chrnb2*, *Th*, *Slc17a6*, *Gad2*) were purchased from Advanced Cell Diagnostics (ACD; http://acdbio.com/). Probes for *Slc17a6*, *Gad2*, and *Th* were previously reported (Wallace et al., 2017; Xiao et al., 2017). Probes for *Chrna4*, *Chrna6*, and *Chrnb2* were validated by comparing co-expression of TH/nAChR subunit mRNA with previously reported analyses using labeled riboprobes (Azam et al., 2002). This prior study indicated that 80-90% of *Th*(+) neurons also expressed *Chrna4*, *Chrna6*, and *Chrnb2*. Consistent with this, we found that 97%, 88%, and 88% of Th(+) neurons in lateral VTA were also positive for *Chrna4*, *Chrna6*, and *Chrnb2*, respectively (**Fig. S1**).

Sections were imaged on a Nikon A1 confocal microscope according to the following parameters: 1024 × 1024 pixels, ~200 nm/pixel, 20X 0.75 NA objective. Nikon system images containing 3 or 4 channels were processed with custom scripts in ImageJ (NIH). All images to be used for FISH quantification were acquired and processed in the same manner. FISH quantification employed the “fluorescence coverage (%)” method (Wallace et al., 2017), which reports the fraction of fluorescent pixels to total pixels in a cellular region of interest (ROI). Multichannel images were opened and all channels (including Dapi staining to identify cellular locations) were overlaid. ROIs were drawn manually around cells containing at least one fluorescent signal, and Dapi staining assisted in distinguishing individual cells from cell clusters. Background subtraction was performed on each channel separately for each image, followed by production of a binary image for quantitative analysis. The percentage of fluorescent pixels to total pixels within each ROI was determined. These data were used to create x vs. y plots of percent coverage for each probe/channel and each cell. To assign a cell as either positive or negative for expression of each probe, a percent coverage “cutoff” was used such that the percent coverage in a ROI had to meet or exceed the cutoff to be counted as positive for the probe. Cutoffs were determined *de novo* for each assay and imaging session to account for any differences in probe performance or tissue autofluorescence over time or across days. A minimum of 3 mice were sampled for each condition, and 3-6 images were analyzed per mouse and per VTA sub-region (midline vs. lateral VTA).

### Brain Slice Preparation and Recording Solutions

Brain slices were prepared as previously described (Engle et al., 2012). Mice were anesthetized with Euthasol (sodium pentobarbital, 100 mg/kg; sodium phenytoin, 12.82 mg/kg) before trans-cardiac perfusion with oxygenated (95% O_2_/5% CO2), 4°C N-methyl-D-glucamine (NMDG)-based recovery solution that contains (in mM): 93 NMDG, 2.5 KCl, 1.2 NaH_2_PO_4_, 30 NaHCO_3_, 20 HEPES, 25 glucose, 5 sodium ascorbate, 2 thiourea, 3 sodium pyruvate, 10 MgSO4-7H_2_O, and 0.5 CaCl_2_H_2_O; 300-310 mOsm; pH 7.3-74). Brains were immediately dissected after the perfusion and held in oxygenated, 4°C recovery solution for one minute before cutting a brain block containing the VTA and sectioning the brain with a vibratome (VT1200S; Leica). Coronal slices (200-250 μm) were sectioned through the VTA and transferred to oxygenated, 33°C recovery solution for 12 min. Slices were then kept in holding solution (containing in mM: 92 NaCl, 2.5 KCl, 1.2 NaH2PO4, 30 NaHCO3, 20 HEPES, 25 glucose, 5 sodium ascorbate, 2 thiourea, 3 sodium pyruvate, 2 MgSO_4_·7H_2_O, and 2 CaCl_2_·2H_2_O; 300-310 mOsm; pH 7.3-74) for 60 min or more before recordings.

Brain slices were transferred to a recording chamber being continuously superfused at a rate of 1.5-2.0 mL/min with oxygenated 32°C recording solution. The recording solution contained (in mM): 124 NaCl, 2.5 KCl, 1.2 NaH_2_PO_4_, 24 NaHCO3, 12.5 glucose, 2 MgSO_4_·7H_2_O, and 2 CaCl_2_·2H_2_O; 300-310 mOsm; pH 7.3-74). For all recordings, the recording solution was supplemented with 1 μM atropine to eliminate contributions from muscarinic ACh receptors.

Patch pipettes were pulled from borosilicate glass capillary tubes (1B150F-4; World Precision Instruments) using a programmable microelectrode puller (P-97; Sutter Instrument). Tip resistance ranged from 5.0 to 10.0 MΩ when filled with internal solution. The following internal solution was used (in mM): 135 potassium gluconate, 5 EGTA, 0.5 CaCl_2_, 2 MgCl_2_, 10 HEPES, 2 MgATP, and 0.1 GTP; pH adjusted to 7.25 with Tris base; osmolarity adjusted to 290 mOsm with sucrose. For uncaging, this internal solution also contained QX-314 (2 mM) for improved voltage control.

### Standard Patch Clamp Electrophysiology

Neurons within brain slices were first visualized with infrared or visible differential interference contrast (DIC), followed in some cases by fluorescence microscopy to identify neurons expressing fluorescent proteins or within range of fluorescent axons. Electrophysiology experiments were conducted using a Nikon Eclipse FN-1 or Scientifica SliceScope. A computer running pCLAMP 10 software was used to acquire whole-cell recordings along with a Multiclamp 700B or Axopatch 200B amplifier and an A/D converter (Digidata 1440A or Digidata 1550A). pClamp software, Multiclamp/Axopatch amplifiers, and Digitata A/D converters were from Molecular Devices. Data were sampled at 10 kHz and low-pass filtered at 1 kHz. Immediately prior to gigaseal formation, the junction potential between the patch pipette and the superfusion medium was nulled. Series resistance was uncompensated. A light emitting diode (LED) light source (XCite 110LED; Excelitas) coupled to excitation filters (400/40 nm, 470/40 nm, and 560/40 nm bandpass) was used to search for fluorescent neurons and, for optogenetic experiments involving ChR2, to stimulate the preparation with light flashes. Light flashes were triggered by pCLAMP via TTL pulses. Flash energy output (0.12 mW/mm^2^) from the LED was determined by calibration using a photodiode power sensor (Model S120C; Thor Labs). To record physiological events following local application of drugs, a drug-filled pipette was moved to within 20−40 μm of the recorded neuron using a second micromanipulator. A Picospritzer (General Valve) dispensed drug (dissolved in recording solution) onto the recorded neuron via a pressure ejection. Ejection volume, duration, and ejection pressure varied depending on the goal of the experiment. Approximate ejection pressures (in p.s.i.) for dispensing saturating ACh concentrations, ACh concentrations for testing galantamine potentiation, and PA-Nic application were 12, 2, and 2, respectively.

### 2-Photon Laser Scanning Microscopy, Electrophysiology, and Nicotine Uncaging

PA-Nic photolysis was performed as previously described (REFERENCE). An Olympus BX51 upright microscope and a 60X (1.0 NA) objective was used to visualize cells. Prairie View 5.4 (Bruker Nano) software was used for image acquisition, photostimulation, and electrophysiology acquisition via a Multiclamp 700B patch clamp amplifier. Analog signals were sampled at 5 kHz and low-pass filtered at 1 kHz, and an A/D converter (PCI-NI6052e; National Instruments) was used for digitization. Patch clamp recordings were carried out using the internal solution mentioned above, except that Alexa 488 or 568 (100 μM) was also included in the recording pipette to visualize cells using 2-photon laser scanning microscopy. After break-in, the internal solution with the Alexa dye was allowed to equilibrate for 15–20 min before imaging was initiated. A Mai Tai HP1040 (Spectra Physics) was used to excite Alexa 488 or 568. Images in **Fig. 1b-d** were acquired by sequentially tuning and imaging, first at 920 nm to image Alexa 488 followed by 1040 nm to image tdT. **Fig. 3n** was acquired by tuning to 1040 nm for tdT imaging. Alexa 568 was used for cell imaging at 790 nm during uncaging experiments. The laser was pulsed at 90 MHz (~250 fs pulse duration), and a M350-80-02-BK Pockels cell (ConOptics) was used for power attenuation. The dual-channel, 2-photon fluorescence was detected by two non-de-scanned detectors; green and red channels (dual emission filters: 525/70 nm and 595/50 nm) were detected by the following Hamamatsu photomultiplier tubes (PMTs), respectively: end-on GaAsP (7422PA-40) and side-on multi-alkali (R3896). A 405 nm continuous wave laser (100 mW OBIS FP LX; Coherent) was used for photostimulation/uncaging via a second set of xy galvanometers incorporated into the scanhead (Cambridge Technologies). 405 nm laser power was measured below the sample but above the condenser using a Field Master GS (LM10 HTD sensor head). PA-Nic (2 mM) was applied locally to the recorded cell via a large-bore (30-40 μm diameter) pressure ejection pipette at low pressure. The Markpoints module of Prairie View 5.4 software was used to select spots in the field of view (~1 μm diameter) for focal uncaging of nicotine via 405 nm laser light flashes (50 ms, 1 mW).

### Statistics and Data Analysis

α level was set to 0.05 for all statistical tests, which were conducted with GraphPad Prism 7 (La Jolla, CA) software. Statistical tests included unpaired students *t*-test, paired *t*-test, and analysis of variance (ANOVA). Error bars are SEM or SD, as indicated in figure legends. Image analysis was performed with ImageJ (NIH). Analysis of electrophysiology data was performed with Clampfit (Molecular Devices) and custom scripts written in MATLAB (The Math Works). Throughout the figure legends, the number of individual neurons tested is stated immediately prior to the number of animals from which those neurons were derived.

## Supplemental Information

**Figure S1. Related to Figure 1.**
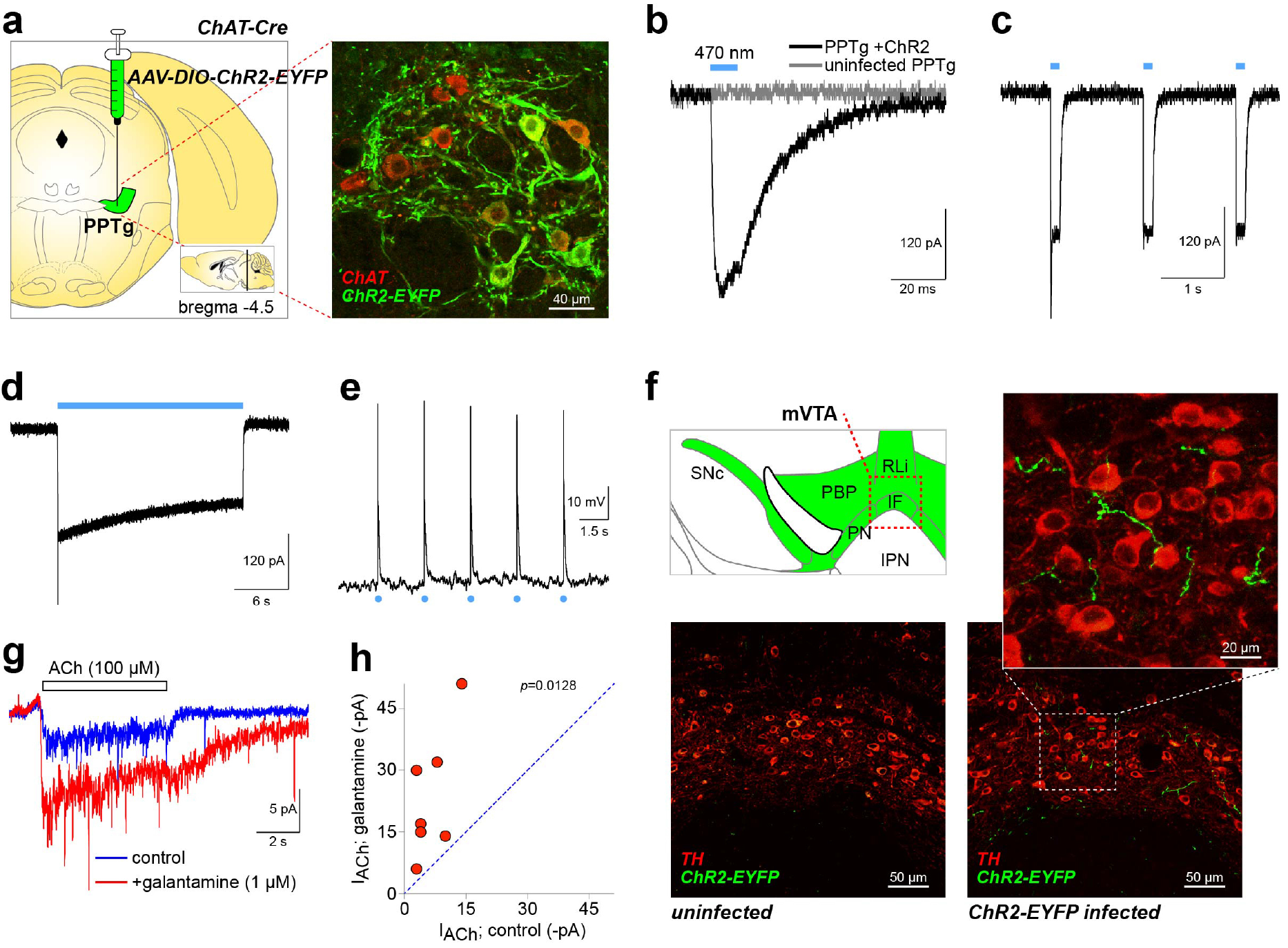
Validation of opsin expression/function in PPTg cholinergic neurons. **(a)** AAVs directing Cre-dependent expression of ChR2-EYFP were bilaterally injected into PPTg of ChAT-Cre mice at the approximate location indicated (left panel). Coronal sections from infected animals were stained with anti-ChAT and anti-GFP antibodies (right panel) to verify ChR2 expression in ChAT neurons. **(b-d)** Photocurrents were recorded in voltage-clamped PPTg neurons expressing ChR2. Brief (10 ms), intermediate (100 ms), and prolonged (20 s) light (470 nm, 0.12 mW/mm^2^) pulses demonstrate ChR2 functionality. **(e)** ChR2 activation drives action potential firing. A ChR2-expressing PPTg ChAT neuron was held in current clamp (I=0) configuration and repetitively stimulated with brief (0.1 ms) light flashes (0.12 mW/mm^2^) to elicit action potentials. **(f)** Expression of ChR2 in mVTA fibers. Coronal sections containing mVTA (top left) from ChAT-Cre mice infected as described in (a) were stained with anti-TH and anti-GFP antibodies to visualize ChR2 expression in cholinergic fibers (top/bottom right). Specificity of detection is provided via uninfected control sections (bottom left). **(g,h)** Galantamine validation. Voltage-clamped mVTA glutamate neurons (n=7/3) were stimulated with pressure ejection application of ACh (100 μM, 5 s, 1-2 psi) to mimic ChR2-mediated ACh release kinetics (**Fig. 1f**). Addition of galantamine potentiated ACh-evoked currents (g). (h) Plot for individual VTA glutamate neurons showing ACh-evoked current amplitude before (control) and after galantamine bath application. *P* value: paired *t*-test.

**Figure S2. Related to Figure 1.**
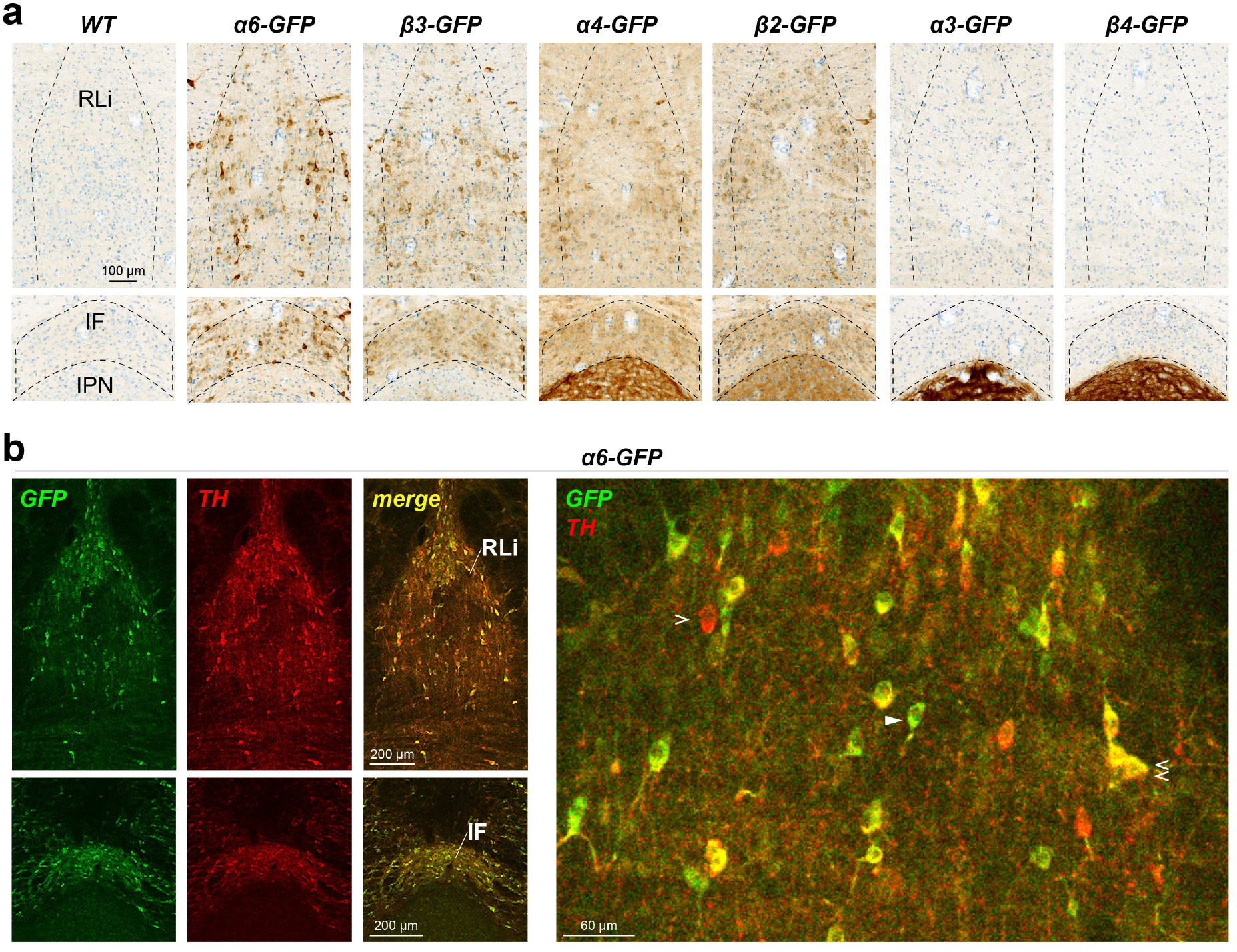
Nicotinic receptor protein expressed in mVTA. **(a)** mVTA nuclei express nAChRs. Coronal sections containing mVTA structures (RLi - rostral linear nucleus; IF - interfascicular nucleus) dorsal to the interpeduncular nucleus (IPN) from knock-in/transgenic mice expressing the indicated GFP-fused nAChR subunit were stained with anti-GFP antibodies and visualized with DAB staining. Representative of n=3 mice. **(b)** The RLi and IF in coronal sections from α6-GFP mice were co-stained with anti-GFP and anti-tyrosine hydroxylase (TH) antibodies (left panels). A high-magnification image through the RLi is shown (right panel), with TH(+)/α6(−) (single open arrowhead), TH(−)/α6(+) (single closed arrowhead), and TH(+)/α6(+) neurons (double open arrowhead).

**Figure S3. Related to Figure 2.**
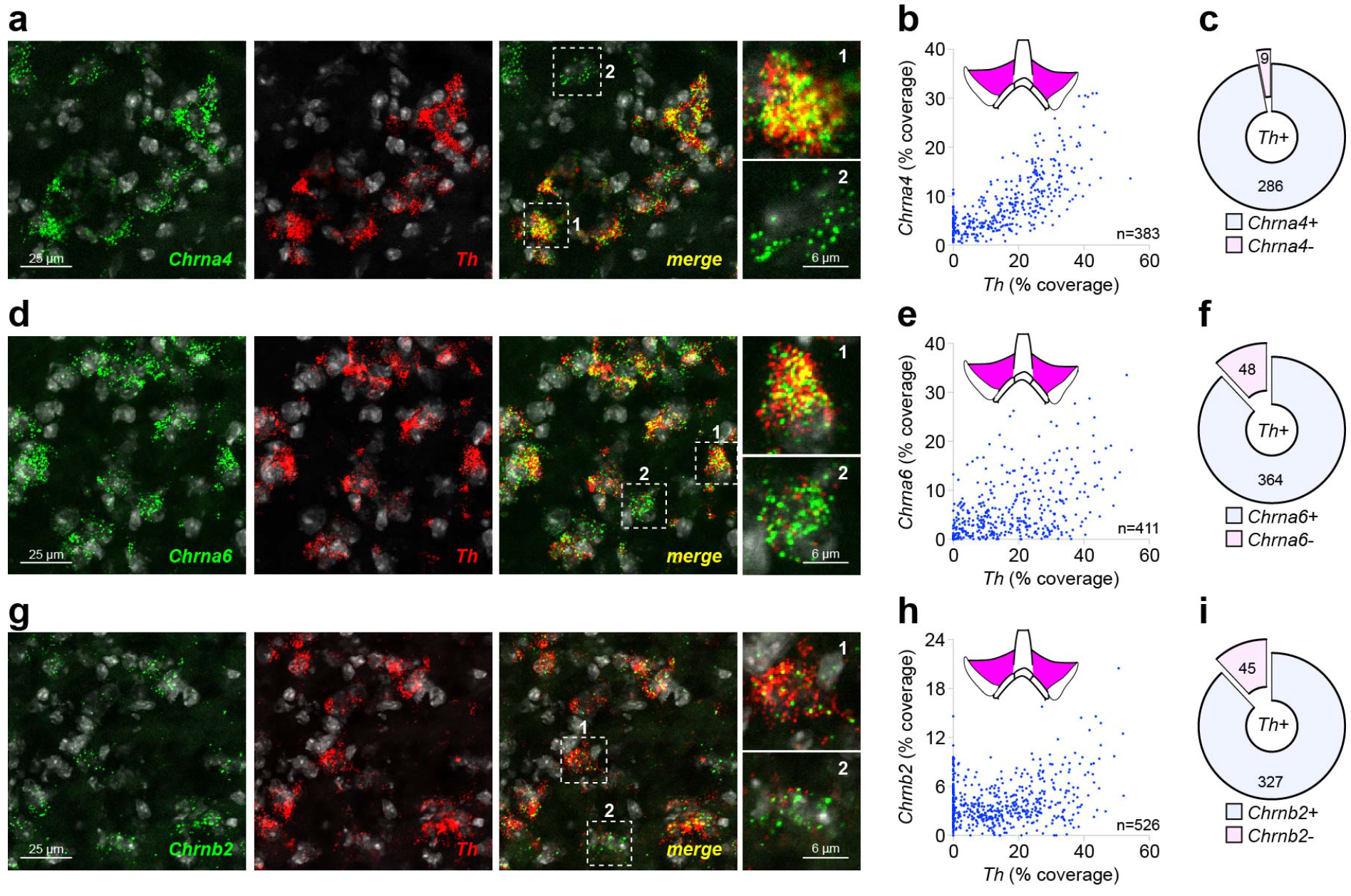
nAChR FISH probe validation. **(a-c)** *Chrna4* FISH probe validation (a) Representative 2-channel fluorescence *in situ* hybridization (FISH) images in lateral VTA neurons for probes: *Chrna4*, *Th*. Single *Chrna4*+ neurons indicated in the ‘merge’ panel, and shown enlarged (right panels), had this expression profile: 1 - *Chrna4*+/*Th*+ 2 - *Chrna4*+/*Th*-. **(b-c)** Analysis of *Chrna4*/*Th* co-expression in lateral VTA. **(b)** Scatter plots showing *Th* (abscissa) and *Chrna4* (ordinate) ‘% coverage’ for all analyzed neurons. **(c)** Pie graph showing the fraction of *Th+* neurons that were *Chrna4*+ vs. *Chrna4*-. **(d-f)** *Chrna6* FISH probe validation. Analysis for *Chrna6* were performed as for *Chrna4*. **(g-i)** *Chrnb2* FISH probe validation. Analysis for *Chrnb2* were performed as for *Chrna4* and *Chrna6*. Data are pooled from n=3 mice.

**Figure S4. Related to Figure 2.**
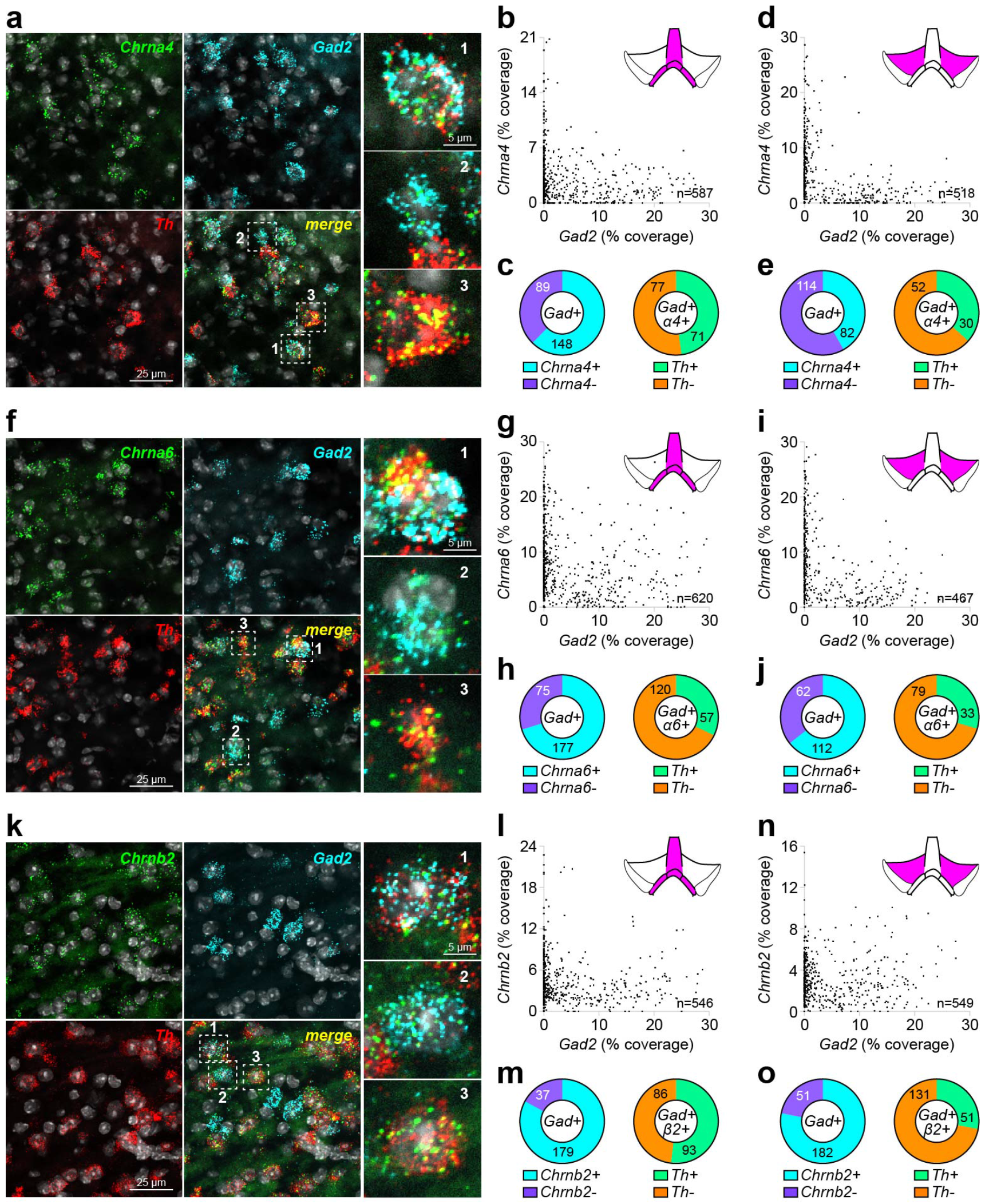
Heteromeric nAChR expression in VTA GABA neurons. **(a-e)** Co-expression of *Chrna4* and *Gad2* in VTA neurons. **(a)** Representative 3-channel fluorescence *in situ* hybridization (FISH) images in mVTA neurons for the following probes: *Chrna4*, *Gad2*, and *Th*. Single *Chrna4*+ neurons indicated in the ‘merge’ panel, and shown enlarged (right panels), had this expression profile: 1 − *Chrna4*+/*Gad2*+/*Th*+; 2 - *Chrna4*+/*Gad2*+/*Th*-; 3 - *Chrna4*+/*Gad2*-/*Th*+. **(b-c)** Analysis of *Chrna4/Gad2* co-expression in mVTA. **(b)** Scatter plots showing *Gad2* (abscissa) and *Chrna4* (ordinate) ‘%coverage’ for all analyzed neurons. **(c)** Left: Pie graph showing the fraction of *Gad2*+ neurons that were *Chrna4*+ vs. *Chrna4*-. Right: *Th* expression status is shown for *Gad2*+/*Chrna4*+ neurons from left graph. **(d-e)** Analysis of *Chrna4/Gad2* co-expression in lateral VTA was performed as in mVTA (b-c). **(f-j)** Co-expression of *Chrna6* and *Gad2* in VTA neurons. Analysis for *Chrna6* were performed as for *Chrna4*. **(k-o)** Co-expression of *Chrnb2* and *Gad2* in VTA neurons. Analysis for *Chrnb2* were performed as for *Chrna4* and *Chrna6*. Complete data for nAChR/*Gad2* FISH is listed in **Supplementary Table 2**. Data are pooled from n=3 mice.

**Figure S5. Related to Figure 4.**
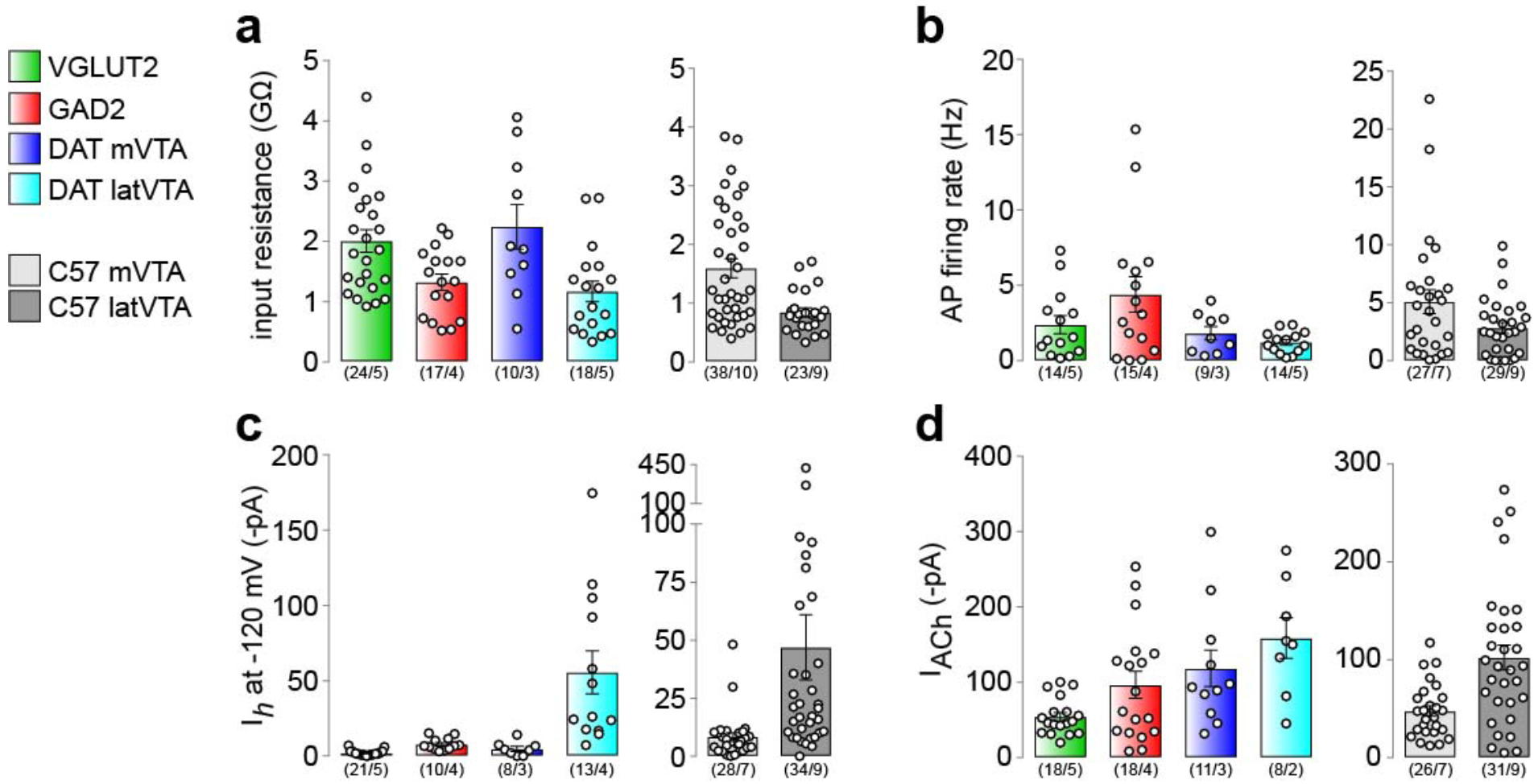
Electrophysiological properties of mVTA neurons. Input resistance **(a)**, action potential firing rate **(b)**, I_*h*_ current at a holding potential of −120 mV **(c)**, and peak current evoked by pressure ejection application of 1 mM ACh **(d)** was measured in mVTA VGLUT2+, GAD2+, and DAT+ neurons in mVTA. DAT+ neurons in lateral VTA were also studied for comparison. The same properties were also measured in medial and lateral VTA neurons in slices from C57Bl/6 WT mice. Number of cells and mice are indicated.

**Supplementary Table 1. Related to Figure 2.**
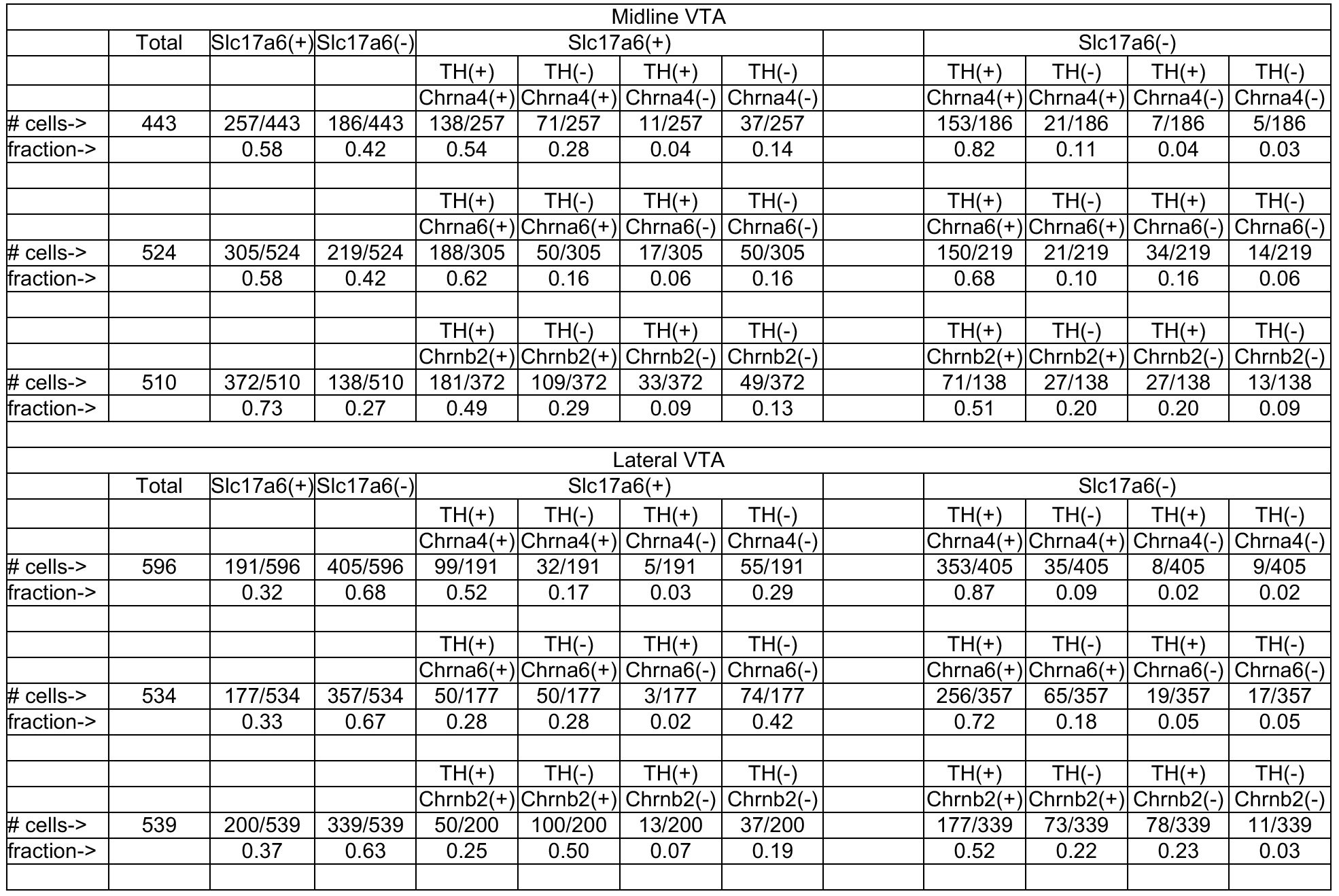
Expression of *Chrna4*, *Chrna6*, and *Chrnb2* in VTA glutamate neurons. Full quantification of triple channel mRNA FISH experiments is shown for midline (top section) and lateral VTA (bottom section). All sections were probed for *Slc17a6* and *Th*, while either *Chrna4*, *Chrna6* or *Chrnb2* was probed in the 3^rd^ channel.

| Midline VTA |  |  |  |  |  |  |  |  |  |  |  |  |
| --- | --- | --- | --- | --- | --- | --- | --- | --- | --- | --- | --- | --- |
|  | Total | Slc17a6(+) | Slc17a6(-) | Slc17a6(+) |  |  |  |  | Slc17a6(-) |  |  |  |
|  |  |  |  | TH(+) | TH(-) | TH(+) | TH(-) |  | TH(+) | TH(-) | TH(+) | TH(-) |
|  |  |  |  | Chrna4(+) | Chrna4(+) | Chrna4(-) | Chrna4(-) |  | Chrna4(+) | Chrna4(+) | Chrna4(-) | Chrna4(-) |
| # cells-> | 443 | 257/443 | 186/443 | 138/257 | 71/257 | 11/257 | 37/257 |  | 153/186 | 21/186 | 7/186 | 5/186 |
| fraction-> |  | 0.58 | 0.42 | 0.54 | 0.28 | 0.04 | 0.14 |  | 0.82 | 0.11 | 0.04 | 0.03 |
|  |  |  |  | TH(+) | TH(-) | TH(+) | TH(-) |  | TH(+) | TH(-) | TH(+) | TH(-) |
|  |  |  |  | Chrna6(+) | Chrna6(+) | Chrna6(-) | Chrna6(-) |  | Chrna6(+) | Chrna6(+) | Chrna6(-) | Chrna6(-) |
| # cells-> | 524 | 305/524 | 219/524 | 188/305 | 50/305 | 17/305 | 50/305 |  | 150/219 | 21/219 | 34/219 | 14/219 |
| fraction-> |  | 0.58 | 0.42 | 0.62 | 0.16 | 0.06 | 0.16 |  | 0.68 | 0.10 | 0.16 | 0.06 |
|  |  |  |  | TH(+) | TH(-) | TH(+) | TH(-) |  | TH(+) | TH(-) | TH(+) | TH(-) |
|  |  |  |  | Chrb2(+) | Chrb2(+) | Chrb2(-) | Chrb2(-) |  | Chrb2(+) | Chrb2(+) | Chrb2(-) | Chrb2(-) |
| # cells-> | 510 | 372/510 | 138/510 | 181/372 | 109/372 | 33/372 | 49/372 |  | 71/138 | 27/138 | 27/138 | 13/138 |
| fraction-> |  | 0.73 | 0.27 | 0.49 | 0.29 | 0.09 | 0.13 |  | 0.51 | 0.20 | 0.20 | 0.09 |
| Lateral VTA |  |  |  |  |  |  |  |  |  |  |  |  |
|  | Total | Slc17a6(+) | Slc17a6(-) | Slc17a6(+) |  |  |  |  | Slc17a6(-) |  |  |  |
|  |  |  |  | TH(+) | TH(-) | TH(+) | TH(-) |  | TH(+) | TH(-) | TH(+) | TH(-) |
|  |  |  |  | Chrna4(+) | Chrna4(+) | Chrna4(-) | Chrna4(-) |  | Chrna4(+) | Chrna4(+) | Chrna4(-) | Chrna4(-) |
| # cells-> | 596 | 191/596 | 405/596 | 99/191 | 32/191 | 5/191 | 55/191 |  | 353/405 | 35/405 | 8/405 | 9/405 |
| fraction-> |  | 0.32 | 0.68 | 0.52 | 0.17 | 0.03 | 0.29 |  | 0.87 | 0.09 | 0.02 | 0.02 |
|  |  |  |  | TH(+) | TH(-) | TH(+) | TH(-) |  | TH(+) | TH(-) | TH(+) | TH(-) |
|  |  |  |  | Chrna6(+) | Chrna6(+) | Chrna6(-) | Chrna6(-) |  | Chrna6(+) | Chrna6(+) | Chrna6(-) | Chrna6(-) |
| # cells-> | 534 | 177/534 | 357/534 | 50/177 | 50/177 | 3/177 | 74/177 |  | 256/357 | 65/357 | 19/357 | 17/357 |
| fraction-> |  | 0.33 | 0.67 | 0.28 | 0.28 | 0.02 | 0.42 |  | 0.72 | 0.18 | 0.05 | 0.05 |
|  |  |  |  | TH(+) | TH(-) | TH(+) | TH(-) |  | TH(+) | TH(-) | TH(+) | TH(-) |
|  |  |  |  | Chrb2(+) | Chrb2(+) | Chrb2(-) | Chrb2(-) |  | Chrb2(+) | Chrb2(+) | Chrb2(-) | Chrb2(-) |
| # cells-> | 539 | 200/539 | 339/539 | 50/200 | 100/200 | 13/200 | 37/200 |  | 177/339 | 73/339 | 78/339 | 11/339 |
| fraction-> |  | 0.37 | 0.63 | 0.25 | 0.50 | 0.07 | 0.19 |  | 0.52 | 0.22 | 0.23 | 0.03 |

**Supplementary Table 2. Related to Figure 2.**
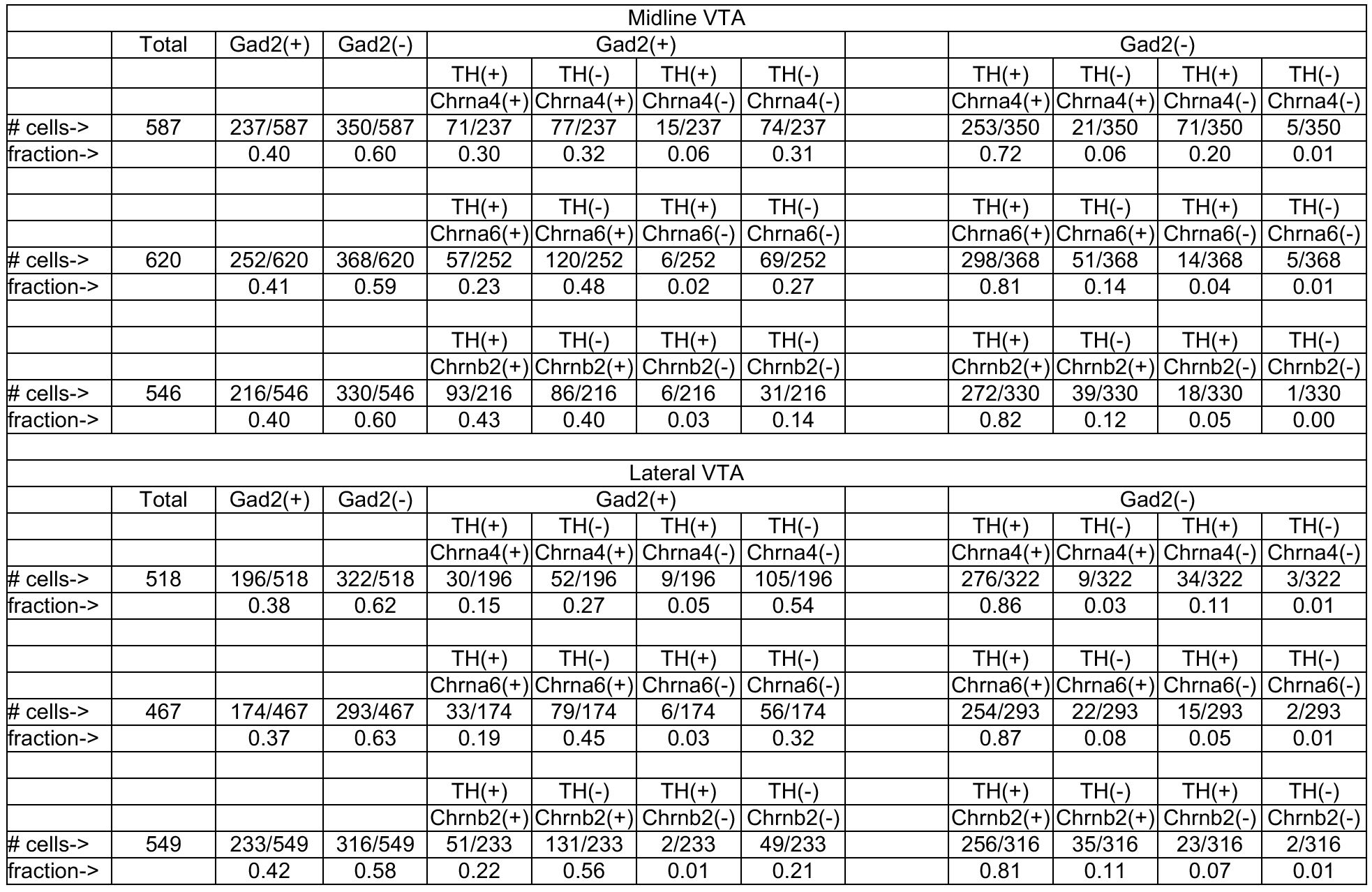
Expression of *Chrna4*, *Chrna6*, and *Chrnb2* in VTA GABA neurons. Full quantification of triple channel mRNA FISH experiments is shown for midline (top section) and lateral VTA (bottom section). All sections were probed for *Gad2* and *Th*, while either *Chrna4*, *Chrna6* or *Chrnb2* was probed in the 3^rd^ channel.

| Midline VTA |  |  |  |  |  |  |  |  |  |  |  |  |
| --- | --- | --- | --- | --- | --- | --- | --- | --- | --- | --- | --- | --- |
|  | Total | Gad2(+) | Gad2(-) | Gad2(+) |  |  |  |  | Gad2(-) |  |  |  |
|  |  |  |  | TH(+) | TH(-) | TH(+) | TH(-) |  | TH(+) | TH(-) | TH(+) | TH(-) |
|  |  |  |  | Chrna4(+) | Chrna4(+) | Chrna4(-) | Chrna4(-) |  | Chrna4(+) | Chrna4(+) | Chrna4(-) | Chrna4(-) |
| # cells-> | 587 | 237/587 | 350/587 | 71/237 | 77/237 | 15/237 | 74/237 |  | 253/350 | 21/350 | 71/350 | 5/350 |
| fraction-> |  | 0.40 | 0.60 | 0.30 | 0.32 | 0.06 | 0.31 |  | 0.72 | 0.06 | 0.20 | 0.01 |
|  |  |  |  | TH(+) | TH(-) | TH(+) | TH(-) |  | TH(+) | TH(-) | TH(+) | TH(-) |
|  |  |  |  | Chrna6(+) | Chrna6(+) | Chrna6(-) | Chrna6(-) |  | Chrna6(+) | Chrna6(+) | Chrna6(-) | Chrna6(-) |
| # cells-> | 620 | 252/620 | 368/620 | 57/252 | 120/252 | 6/252 | 69/252 |  | 298/368 | 51/368 | 14/368 | 5/368 |
| fraction-> |  | 0.41 | 0.59 | 0.23 | 0.48 | 0.02 | 0.27 |  | 0.81 | 0.14 | 0.04 | 0.01 |
|  |  |  |  | TH(+) | TH(-) | TH(+) | TH(-) |  | TH(+) | TH(-) | TH(+) | TH(-) |
|  |  |  |  | Chrb2(+) | Chrb2(+) | Chrb2(-) | Chrb2(-) |  | Chrb2(+) | Chrb2(+) | Chrb2(-) | Chrb2(-) |
| # cells-> | 546 | 216/546 | 330/546 | 93/216 | 86/216 | 6/216 | 31/216 |  | 272/330 | 39/330 | 18/330 | 1/330 |
| fraction-> |  | 0.40 | 0.60 | 0.43 | 0.40 | 0.03 | 0.14 |  | 0.82 | 0.12 | 0.05 | 0.00 |
| Lateral VTA |  |  |  |  |  |  |  |  |  |  |  |  |
|  | Total | Gad2(+) | Gad2(-) | Gad2(+) |  |  |  |  | Gad2(-) |  |  |  |
|  |  |  |  | TH(+) | TH(-) | TH(+) | TH(-) |  | TH(+) | TH(-) | TH(+) | TH(-) |
|  |  |  |  | Chrna4(+) | Chrna4(+) | Chrna4(-) | Chrna4(-) |  | Chrna4(+) | Chrna4(+) | Chrna4(-) | Chrna4(-) |
| # cells-> | 518 | 196/518 | 322/518 | 30/196 | 52/196 | 9/196 | 105/196 |  | 276/322 | 9/322 | 34/322 | 3/322 |
| fraction-> |  | 0.38 | 0.62 | 0.15 | 0.27 | 0.05 | 0.54 |  | 0.86 | 0.03 | 0.11 | 0.01 |
|  |  |  |  | TH(+) | TH(-) | TH(+) | TH(-) |  | TH(+) | TH(-) | TH(+) | TH(-) |
|  |  |  |  | Chrna6(+) | Chrna6(+) | Chrna6(-) | Chrna6(-) |  | Chrna6(+) | Chrna6(+) | Chrna6(-) | Chrna6(-) |
| # cells-> | 467 | 174/467 | 293/467 | 33/174 | 79/174 | 6/174 | 56/174 |  | 254/293 | 22/293 | 15/293 | 2/293 |
| fraction-> |  | 0.37 | 0.63 | 0.19 | 0.45 | 0.03 | 0.32 |  | 0.87 | 0.08 | 0.05 | 0.01 |
|  |  |  |  | TH(+) | TH(-) | TH(+) | TH(-) |  | TH(+) | TH(-) | TH(+) | TH(-) |
|  |  |  |  | Chrb2(+) | Chrb2(+) | Chrb2(-) | Chrb2(-) |  | Chrb2(+) | Chrb2(+) | Chrb2(-) | Chrb2(-) |
| # cells-> | 549 | 233/549 | 316/549 | 51/233 | 131/233 | 2/233 | 49/233 |  | 256/316 | 35/316 | 23/316 | 2/316 |
| fraction-> |  | 0.42 | 0.58 | 0.22 | 0.56 | 0.01 | 0.21 |  | 0.81 | 0.11 | 0.07 | 0.01 |

